# Fluctuating landscapes and heavy tails in animal behavior

**DOI:** 10.1101/2023.01.03.522580

**Authors:** Antonio Carlos Costa, Massimo Vergassola

## Abstract

Animal behavior is shaped by a myriad of mechanisms acting on a wide range of scales. This immense variability hampers quantitative reasoning and renders the identification of universal principles elusive. Through data analysis and theory, we here show that slow non-ergodic drives generally give rise to heavy-tailed statistics in behaving animals. We leverage high-resolution recordings of *C. elegans* locomotion to extract a self-consistent reduced order model for an inferred reaction coordinate, bridging from sub-second chaotic dynamics to long-lived stochastic transitions among metastable states. The slow mode dynamics exhibits heavy-tailed first passage time distributions and correlation functions, and we show that such heavy tails can be explained by dynamics on a time-dependent potential landscape. Inspired by these results, we introduce a generic model in which we separate faster mixing modes that evolve on a quasi-stationary potential, from slower non-ergodic modes that drive the potential landscape, and reflect slowly varying internal states. We show that, even for simple potential landscapes, heavy tails emerge when barrier heights fluctuate slowly and strongly enough. In particular, the distribution of first passage times and the correlation function can asymptote to a power law, with related exponents that depend on the strength and nature of the fluctuations. We support our theoretical findings through direct numerical simulations.

## I. INTRODUCTION

Throughout their lives, animals continuously sense, process information and act appropriately to ensure survival. Such ever-changing behavior emerges from orchestrated biological activity across a wide range of scales. The intrinsic high-dimensionality of these far from equilibrium dynamics, where multiple timescales continuously interact, pose deep challenges to a quantitative understanding. Yet, time is ripe for theory. Recent advances in machine vision technologies (e.g., [1–3]) make it possible to record an animal’s pose in unconstrained environments with unprecedented resolution. Such data now span several orders of magnitude in space and time [4], challenging our quantitative understanding and demanding modeling approaches that can bridge across scales: from sub-second movements to hours-long strategies.

Despite these technical advances, we are still far from measuring all the variables that determine behavior. Indeed, a complete microscopic theory would require knowledge not only of the current posture of the animal, but also its physiological state, its sensors, the state of its muscle cells, the kinetics of an uncountable number of molecules that determine the interaction between neurons, the genetic expression of different types of neuromodulators, and so on. However, examples from statistical physics highlight how microscopic details might not be required to predict the dynamics of carefully selected collective variables [5]. Indeed, to study how odor molecules diffuse in the air we need not measure the position and momentum of each molecule. Instead, we can simply write down a self-determined equation for the transport of the concentration field itself [6]. In fact, much of the success of statistical mechanics relies on the identification of slowly varying macroscopic modes, which, through a time-scale separation, depend only statistically on the microscopic details of the dynamics. Finding such order parameters, however, is typically not a simple task, and requires a build-up of intuition that is often not accessible for far-from-equilibrium systems as encountered in biology. Nonetheless, we leverage the notion of time scale separation to search for such slowly-varying collective variables and show that it is possible to build tractable reduced-order models directly from imaging data of behaving animals. Additionally, through a combination of data analysis and theory, we show that the ever-changing nature of biological systems, which evolve at a wide range of time scales, provides a minimal yet sufficient mechanism for generating heavy-tailed statistics.

Our inspiration is the foraging behavior of the nematode C. elegans, a pivotal model organism in genetics and neuroscience [7, 8]. On a two-dimensional agar plate, worms move by propagating dorsoventral waves throughout their bodies; on short time scales, they control the frequency, wavelength, and direction of those waves to move forward, backward, or turn. Sequences of such short-lived movements exhibit signatures of chaos generating temporal variability in the worm’s behavior [9, 10]. Despite this inherent unpredictability, Costa et al. [11] showed that by reconstructing unobserved influences through a time-delay embedding [12–16], it is possible to build a high-fidelity Markov model that accurately predicts *C. elegans* foraging behavior, rendering simulated worms nearly indistinguishable from real worms across a wide range of scales. This Markov model also recovered coarse-grained descriptions of the worm’s foraging strategy, identifying long-lived metastable states that correspond to transitions between relatively straight paths (“runs”) and not-so-abrupt reorientations (“pirouettes”) [11, 17] (a two-state characterization that is akin to the run-and-tumbling of bacteria [18, 19]). Recent analysis of posture-scale movements highlights how other organisms also exhibit stereotyped movements [20–22], making stereotypy one of the few general *principles* in the physics of animal behavior [4, 23].

The emergence of stereotypy from continuous movement stems from an implicit time scale separation between variations on what is defined as a behavioral state, and transitions between behavioral states, much like particles hopping among wells in potential landscapes. Here, we make this evocative picture concrete. In the first section, we infer an effective Langevin description for the worm’s “run-and-pirouette” dynamics. Notably, we find long-range correlations and heavy-tailed distributions of times spent either performing a “run” or a “pirouette”, instead of the exponential timescales expected for independent transition events with a fixed hopping rate. We then show how such nontrivial statistics stem from the slow adaptation of the worm’s search strategy, which is captured by time-dependent model parameters that we infer directly from the data. In the second section, we investigate whether non-ergodic fluctuations (such as the worm’s adaptation) generally give rise to heavy-tailed statistics. We introduce a generic model of animal behavior in which the posture dynamics evolves in potential landscapes that fluctuate in time, and show that even simple potential landscapes can exhibit heavy-tailed first passage times and long-range correlations when barrier heights fluctuate in time slowly and strongly enough.

## II. HEAVY TAILS IN *C. ELEGANS* BEHAVIOR: THE ROLE OF ADAPTATION

We leverage a previously analyzed dataset in which 12 lab-strain N2 worms are placed on an agar plate and allowed to freely explore for 35 minutes [24] (see Appendix A). From each video frame (sampled every *δt* = 1*/*16 s), we extract the worm’s centerline, measure tangent angles equally spaced along the body, and subtract the overall rotation of the worm to obtain a worm-centric representation of the animal’s shape *θ*_*t*_. As done in [9, 11], we then stack *K*^∗^ = 11 time delays of the animals posture *X* _*K*∗_ = *θ*_*t:t*+*K*∗_ to obtain a maximally-predictive sequence space, Fig. 1(a). In this way, we subsume the short-term memory resulting from *hidden* dynamics into an expanded state space that admits an approximately Markovian description. Assuming stationary dynamics and a fully resolved state, the dynamics is then given by

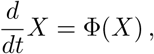

where Φ(*X*) is a nonlinear noisy function that evolves the state *X*_*K*∗_. The corresponding evolution of probability densities *ρ*_*t*_ = *ρ*(*X*_*K*∗_, *t*) is given by,

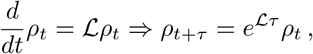

where Φ is encoded into a linear transfer operator ℒ that evolves probability densities. Given an appropriate discretization of the state space, it is possible to approximate the action of ℒ as a Markov chain [10, 11].

**FIG. 1.**
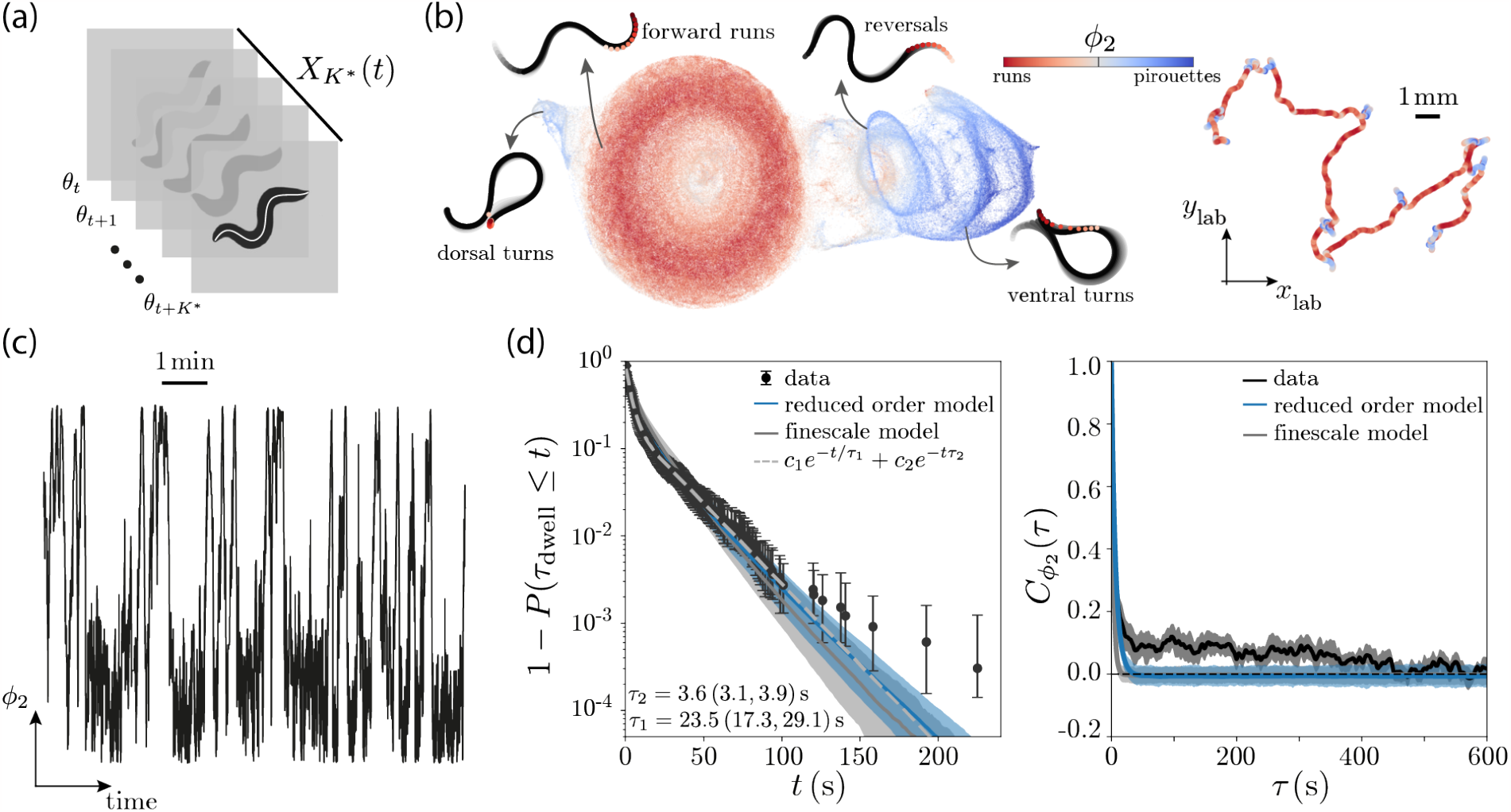
A reduced-order model of *C. elegans* foraging dynamics. (a) From video imaging data, we extract the body posture in a worm-centric perspective by measuring local tangent angles along the body and rotating them onto a common frame-of-reference, obtaining *θ*_*t*_ [26]. These vectors are then stacked over a timescale *K*^∗^ to yield maximally predictive states *X* _*K*∗_ (*t*) [9–11]. (b) Through an appropriate fine-scale partitioning of *X* _*K*∗_ (*t*), we obtain a high-fidelity Markov model of the dynamics [10, 11]. The eigenspectrum of the inferred Markov chain captures a hierarchy of timescales, with the first non-trivial eigenvector of the reversibilized transition matrix *ϕ*_2_ capturing transitions between “runs” and “pirouettes”. We represent the high-dimensional state-space *X* _*K*∗_ through a 2D UMAP embedding as in [11] (left), and color-code each point in this space by the projection along *ϕ*_2_ (see Appendix A). Each point in this space corresponds to a *K*^∗^ sequence of postures and different behaviors correspond to different regions of the space. We also plot an example 10 min long centroid trajectory color-coded by *ϕ*_2_ (right). The example states and the centroid trajectory showcases how *ϕ*_2_ *<* 0 corresponds to forward “runs” whereas *ϕ*_2_ *>* 0 corresponds to combinations of reversals, ventral and dorsal turns used during “pirouettes”. (c) Example time series of *ϕ*_2_ illustrating the stochastic hopping between “runs” and “pirouettes”. (d-left) First passage time distribution obtained from the data (black), simulations of the stochastic dynamics of Eq. 1 (blue) and simulations performed with the full model ℒ (gray). The bulk of the distribution is captured by a sum of exponential functions (gray dashed line) that are predicted by the simulations, but the data also exhibits heavy tails that are not captured accurately. (d-right) Connected autocorrelation function *C*_*ϕ*2_ (*τ*) for the data (black) and simulations performed with the reduced-order model of Eq. 1 (blue) and the full model ℒ (gray). While model simulations capture the dynamics over short timescales, they fail to predict the long-range correlations exhibited by the data. Notably, the stochastic model of Eq. 1 gives predictions for the first passage time distribution and autocorrelation functions that are comparable with those obtained with the full model ℒ [11], showcasing the self-consistency of this model at capturing the long-lived dynamics. Error bars represent 95% confidence intervals bootstrapped across worms.

Encoding the nonlinear dynamics of *X*_*K*∗_ into the linear evolution of the probability distribution, offers a means to a principled coarse-graining. Indeed, the eigen-decomposition of ℒ, 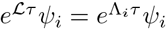, yields a set of eigen-values that capture the hierarchy of timescales by which different eigenfunctions relax to the steady-state (Λ_*i*_ has units of inverse time). For a mixing system, there is a unique largest eigenvalue Λ_1_ = 0 that corresponds to the steady-state distribution *ψ*_1_ = *π* = *e*^0*τ*^ *π*. The remaining eigenfunctions *ψ*_*i>*1_, organized according to increasing real parts, correspond to collective variables that relax to the steady-state on faster and faster timescales, set by |Re(Λ_*i>*1_) |. For *C. elegans* foraging dynamics, the eigenspectrum of the reversibilized ℒ reveals a main slow mode *ϕ*_2_, which was used to coarse-grain the behavior into “runs” and “pirouettes” [11]. Here, we leverage this timescale-separated eigenfunction to define a slow reaction coordinate [25] that captures transitions along a “run-and-pirouette” axis (see Appendix A), Fig. 1(b). In particular, we project the full posture dynamics onto *ϕ*_2_, going from the fast chaotic dynamics of the body posture [9] to an effective stochastic description for the hopping between “runs” and “pirouettes”. An example time series of *ϕ*_2_(*t*) is shown in Fig. 1(c).

### A. Inferring a stationary Langevin equation for the “run-and-pirouette” dynamics

We here aim to infer an explicit coarse-grained model for the apparent stochastic hopping along *ϕ*_2_(*t*). Projecting the full dynamics onto *ϕ*_2_, however, results in non-Markovian effects due to the fact that the orthogonal projection (the modes we do not take into account) includes non-vanishing temporal correlations coming from the faster-decaying eigenfunctions of ℒ [27–29]. To deal with this, we sample the dynamics on a timescale *τ*^∗^ long enough such that the temporal correlations in the noise have decayed [30]. In this way, we can obtain an effective overdamped Langevin description for *ϕ*_2_(*t*) [31],

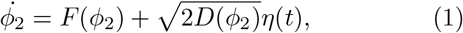

in which, since the dynamics are sampled every *τ*^∗^, we effectively have ⟨*η*(*t*)*η*(*t*^*′*^) ⟩ = *δ*(*t* −*t*^*′*^), *t* = *nτ*^∗^, *t*′ = *n*′*τ*^∗^, *n, n*′ ∈ℕ. Indeed, with *τ*^∗^ = 0.75 s the slow dynamics becomes effectively Markovian [11], and a stochastic model inferred from the time series results in effectively delta-correlated fluctuations, Fig. S1(b). To find *F* (*ϕ*_2_) and *D*(*ϕ*_2_) we leverage a kernel-based approach [32], based on the Kramers-Moyal expansion (e.g., [33]). Instead of estimating the drift and diffusion coefficients in discretized bins, we use kernels to obtain a more robust and continuous estimate of *F* (*ϕ*_2_) and *D*(*ϕ*_2_) (see Appendix A).

To probe the ability of this model to reproduce the long-lived properties of the dynamics, we identify “run” and “pirouettte” states by maximizing the metastability of both states (see Appendix A) [11], and estimate the time spent in these two behaviors, Fig. 1(d-left), Fig. S2. Interestingly, while the main exponential time scales of the first passage time distribution (FPTD) are captured by the inferred stationary Langevin dynamics, the data also exhibits a heavier tail that this model naturally does not predict. In addition, we estimated the connected autocorrelation function

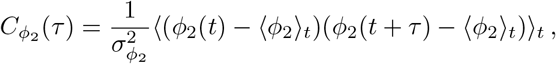

Where 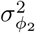 is the variance of *ϕ*_2_(*t*) and ⟨· ⟩_*t*_ represents a temporal average. In accordance with the first passage time results, we observe that, while the model captures correlations on relatively short timescales (≈10 s), the data exhibits long-range correlations that the model fails to predict, Fig. 1(d-right). While this discrepancy could be associated with the projection onto a single slow mode, or with the assumption of Langevin dynamics, we find that simulating the dynamics using the full model ℒ [11] results in similar predictions, showcasing the self-consistency of our coarse-graining approach, Fig. 1(d).

### B. Fluctuating potential landscapes underlie the emergence of heavy tails in *C. elegans* foraging

While the inferred ℒ captures *C. elegans* foraging behavior across several scales [11], it fails to predict the heavy-tailed statistics observed at the longest times. One possible explanation for the inability of the Langevin dynamics of Eq. 1 (or the full model ℒ) to capture these observations, is the existence of subtle *hidden* fluctuations that evolve on timescales comparable to the observation time and are not accurately captured by our time-delay embedding, rendering the dynamics non-stationary. Indeed, it has been observed that upon removal from food worms slowly adapt their search strategy by lowering their rate of “pirouettes” to explore wider areas in search for food [34–39]. In order to have a time-evolving rate of pirouettes we would need to extend the stationary model to allow for explicitly time-dependent drift and diffusion terms,

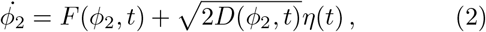

that reflect the adaptation of the worm’s foraging strategy throughout the experimental time scales.

We infer time-dependent drift and diffusion coefficients in overlapping windows, defined to be long enough to equilibrate the fast dynamics but short enough such that the steady-state distribution does not change significantly (see Appendix A). Interestingly, the time evolution of the effective potential landscape, Fig. 2(a), shows that worms slowly adapt their search strategy by increasingly performing runs, in agreement with previous studies [34–39]. Over time, the “run-and-pirouette” random walk is therefore biased to explore further away in search of food. Notably, this explicitly time-dependent model is sufficient to quantitatively reproduce the heavy tail of the first passage time distribution and the nontrivial long-range correlations exhibited by real worms, Fig. 2(b).

**FIG. 2.**
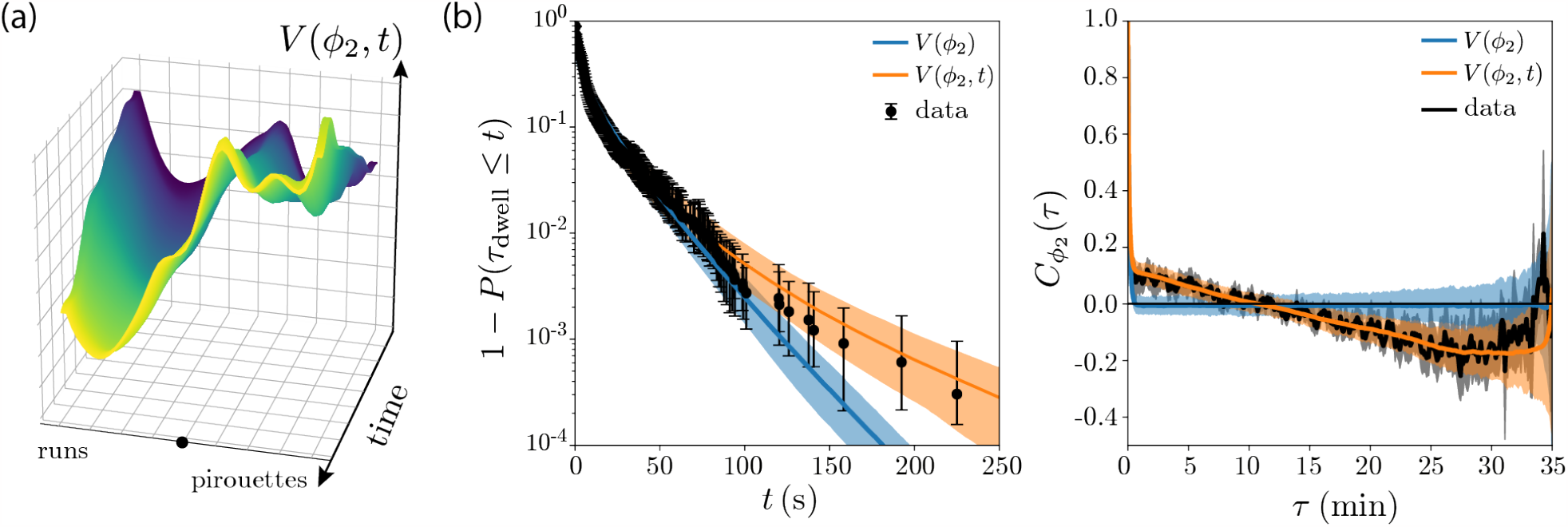
A time-varying potential landscape captures heavy tails in *C. elegans* foraging behavior. (a) We infer a time-dependent potential landscape description from the *ϕ*_2_ time series by estimating drift and diffusion coefficients in sliding windows (see Appendix A), and show the result for an example worm. Notably, as time goes on (blue to yellow) we observe a biasing of the behavior towards increasingly performing “runs”. (b) First passage time distribution (left) and connected autocorrelation function 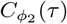 (right) obtained from the data (black), simulations from the static model of Eq. 1 (blue, same as Fig. 1(d)) and simulations using a time-dependent potential landscape, Eq. 2 (orange). Notably, including an explicit time-dependence to the model parameters recovers the heavy tailed first passage times and the non-trivial long-range correlations observed in the data. Error bars represent 95% confidence intervals bootstrapped across worms.

## III. LONG TIMESCALES AND THE EMERGENCE OF HEAVY TAILS IN ANIMAL BEHAVIOR

These results show that the observed heavy tails result from the adaptation of the worms’ search strategy. Could similar mechanisms underlie the widespread observation of heavy tails across behaving animals? Animals do modulate their behavior across a vast range of scales, either due to environmental factors or through endogenous fluctuating internal states driven by neuromodulation, such as hunger or stress [40–42]. Such a continuum of scales would inevitably result in non-stationary dynamics since long-lived modes would prevent the relaxation to a steady-state distribution within a finite observation time *T*_exp_. We here investigate if such non-stationary fluctuations give rise to heavy-tailed statistics. In particular, we introduce a general picture of behavior in which the pose dynamics evolves in potential landscapes that fluctuate over time.

### A. A fluctuating landscape picture of animal behavior

Given a set of observations of animal locomotion (e.g., from video imaging), we consider that the dynamics can be decomposed into ergodic, *x*, and non-ergodic, *s*, components. The former are the state-space variables that mix sufficiently well and define the potential wells that correspond to the stereotyped behaviors ; the latter evolve on time scales *τ*_*s*_ comparable to the observation time and slowly modulate the potential landscape of *x*. Assuming that we can simplify the dynamics onto an overdamped Langevin description through an appropriate time scale separation, we obtain a phenomenological model of the long-lived dynamics as a system of Itô stochastic differential equations,

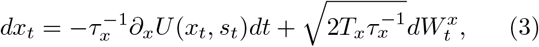

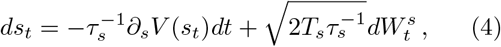

where we set *τ*_*x*_ = 1 without loss of generality (what matters is the ratio *τ*_*s*_*/τ*_*x*_), 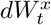 and 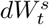 are independent increments of a Wiener process, *T*_*x*_ and *T*_*s*_ capture the level of fluctuations in *x* and *s* respectively, *U* is a potential landscape where different wells correspond to long-lived stereotyped behaviors and *V* is assumed to be uncoupled from the dynamics of *x* for simplicity. In the following sections, we will that the slow modulation of the barrier heights of *U* (*x, s*) through the dynamics of *s* is sufficient to explain the emergence of heavy-tailed first passage time distributions and the non-trivial correlations we observed for the worm behavior, Fig. 2(b).

### B. Heavy-tailed first passage times in slowly-driven metastable dynamics

In the context of the Langevin dynamics of Eq. 3, the distribution of times spent in a given behavioral state is given by the statistics of escape from a potential well, a well-studied problem in statistical physics [43]. Despite the general interest in this concept across several fields [44–47], finding analytical expressions for the density of first passage time events is generally a formidable task [48]. Indeed, most results focus on the mean first passage time (MFPT), which is more tractable (see, e.g., [49, 50]). However, the MFPT provides only limited information: when multiple time scales are involved, the MFPT is not representative of the long time behavior of the distribution [51]. To investigate whether the non-ergodic dynamics of Eqs. 3,4 can give rise to heavy tails, we here derive the large time asymptotic behavior of the first passage time distribution.

The measurement time *T*_exp_ separates ergodic from non-ergodic dynamics. Importantly, it also sets a lower bound on the slowest observed hopping rates 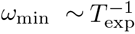, such that when *τ*_*s*_ = 𝒪 (*T*_exp_) we can make an adiabatic approximation and assume that transition events occur within a nearly static potential. The long-time behavior of the first passage time distribution is dominated by the deepest potential well, which when stationary yields a first passage time distribution

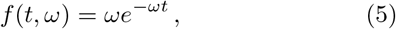

where *ω* is the dominating slow kinetic transition rate that we assume to be dependent on *s*. When we allow *s* to fluctuate, *ω* also varies, and the distribution of first passage times *f* (*t*) is given by the expectation value of *f* (*t, ω*) over the distribution of *ω, p*(*ω*), weighted by the effective number of transition observed within *T*_exp_, which is proportional to *ω*. Marginalizing over *ω* we get

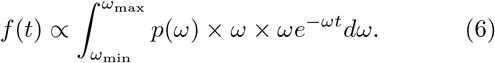

While the barrier height depends on the dynamics of a slow control parameter *s*, the tail of the distribution is dominated by instances in which the barrier height is the largest, motivating the use of Kramers approximation (see, e.g., [43, 52]),

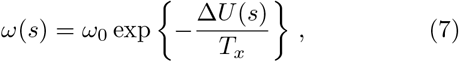

where ∆*U* (*s*) = *U* (*x*_*f*_, *s*) − *U* (*x*_0_, *s*) and *ω*_0_ is a constant (see Appendix B). Assuming that each measurement starts from different initial conditions sampled according to a Boltzmann weight, the distribution of *s* is given by [53],

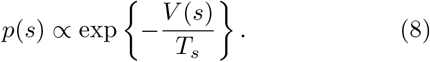

As shown in Appendix B, when the barrier height fluctuations are large enough to yield an 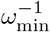 that is comparable to the measurement time 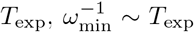, we can combine equations Eqs. 6, 7,8 to obtain an asymptotic approximation of the FPTD in the large *t* limit,

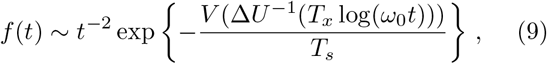

where ∆*U*^−1^(·) represents the inverse function of ∆*U* (*s*) and we have kept only the dominant order of the asymptotic approximation (see Appendix B). Importantly, when *T*_*s*_ we obtain *f* (*t*) ∼ *t*^−2^ under very general assumptions for the form of *V* (*s*) and *U* (*x, s*). In addition, when *V* (*s*) and ∆*U* (*s*) are asymptotically equivalent, *f* (*t*) behaves as a power law 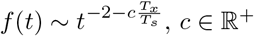 with an exponent that depends on the ratio between the fluctuations in *x* and in *s*. This derivation qualitatively recovers what we observed for the worm behavior, Fig. 2, showcasing how slow non-ergodic drives can indeed give rise to heavy-tailed first passage time distributions. In the following section, we demonstrate these results with illustrative examples.

#### 1. Poisson process with varying hopping rates

To probe our theoretical predictions, we first assume that hopping events are well captured by a Poisson process and that the modulation of the potential landscape is infinitely slow such that the adiabatic approximation of Eq. 6 holds exactly. In practice, we sample *s* according to the Boltzmann distribution, Eq. 8, and in order to relax from the Kramers approximation, we obtain *ω*(*s*) through the backward Kolmogorov equation [52],

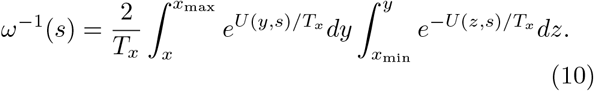

We then sample events according to the distribution of first passage times *f* (*t, ω*), Eq. 5, until reaching the measurement time *T*_exp_, Fig. 3(a). We take *U* (*x, s*) = *s*^2^(*x*^2^ − 1)^2^ to be a symmetric double well potential and sample *s* according to a Boltzmann distribution with 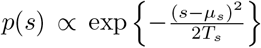, corresponding to an Ornstein-Uhlenbeck process. Since the first passage time distribution is dominated by the large energy barriers, *s* ≫ *μ*_*s*_, we take *V* (∆*U*^−1^(*x*)) ∼ *x/*2. From the derivation of Eq. 9, we expect that the final distribution of first passage times will be given by,

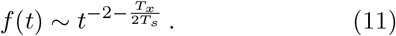

**FIG. 3.**
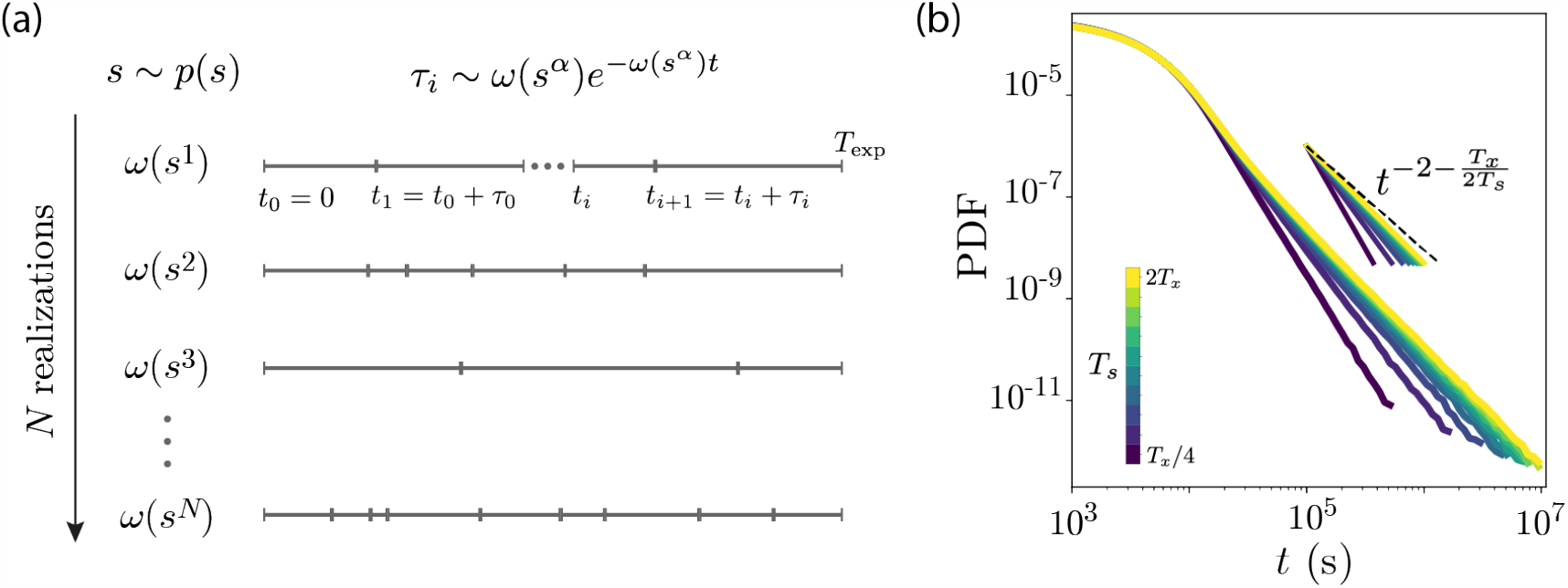
Heavy-tailed first passage time distribution in Poisson process with varying hopping rates. (a) Schematic of the simulation process. For each realization, we sample *s* according to the Boltzmann distribution, *p*(*s*). The hopping rate corresponding to a particular sample *s*^*i*^ is then determined by the backward Kolmogorov equation, Eq. 10, and event durations are sampled according to the first passage time distribution *f* (*t, ω*) = *ωe*^−*ωt*^ until reaching the experimental timescale *T*_exp_. This process is then repeated over *N* =50,000 realizations (see Appendix A). (b) First passage time distribution for the Poisson process with varying hopping rates. We collect the duration of the event from our simulations and estimate their probability distribution function (PDF). As predicted, we obtain a power law with an exponent 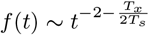.

Indeed, this is what we find through numerical simulations, Fig. 3(b).

#### 2. Slowly-driven double-well potential

We now relax the simplifications of the previous analysis and fully simulate the two-dimensional stochastic dynamics for a double-well potential whose barrier height is slowly modulated according to an Ornstein-Uhlenbeck process, Fig. 4(a). The dynamics are given by an Itô stochastic differential equation,

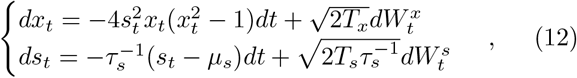

where *T*_*x*_ = 10^−3^, 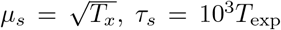, *τ*_*s*_ = 10^3^*T*_exp_ and 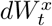, 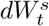 are independent increments of a Wiener process (see Appendix A for simulation details). Since the tail of *f* (*t*) is dominated by large *s* values, we again take *V* (*s*) ∼ *s*^2^*/*2, and thus *V* (∆*U*^−1^(*x*)) ∼ *x/*2. From the derivation of Eq. 9 we find,

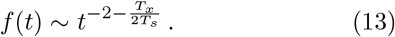

**FIG. 4.**
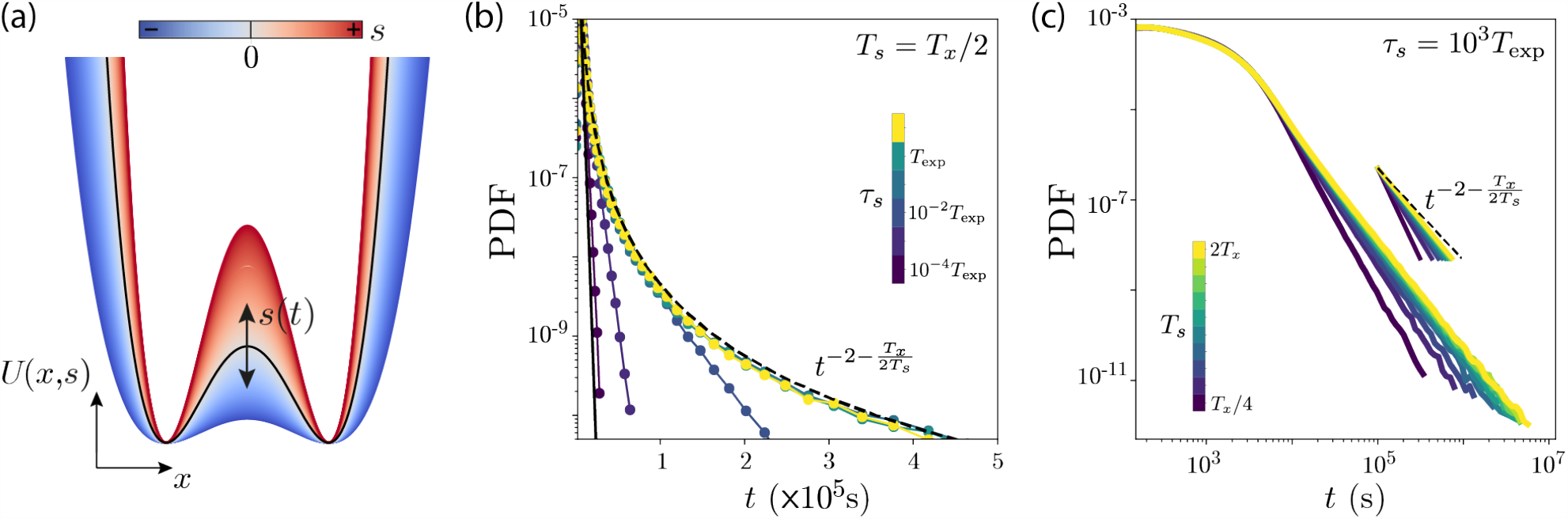
Emergence of heavy-tails in the first passage time distribution of a slowly-driven double-well potential. Schematic of the variation in the double-well potential with *s* (colored from blue to red; the black line represents *s* = *μ*_*s*_). Probability distribution function (PDF) of the first passage times obtained from direct numerical simulations of Eq. 12 for different values of *τ*_*s*_ and *T*_*s*_ = *T*_*x*_*/*2 (see Appendix A). When *τ*_*s*_ → 0, the potential landscape relaxes to its mean value much faster than the time it takes to escape the well, resulting in exponential behavior with a hopping rate corresponding to *μ*_*s*_ (black line). As *τ*_*s*_ approaches *T*_exp_, we start observing a transition from exponential to power law behavior, and in the limit of large *τ*_*s*_ we obtain the power law behavior derived in Eq. 13 (black dashed line). (c) Estimated FPTD from direct numerical simulations of Eq. 12 for large *τ*_*s*_ = 10^3^*T*_exp_ and different values of *T*_*s*_ (see Appendix A). As predicted, the tail of the distribution behaves as a power law 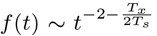 (colored lines) with an exponent that approaches −2 as *T*_s_ → ∞ (black dashed line).

To test this result we performed direct numerical simulations of Eq. 12 while varying *T*_*s*_ and *τ*_*s*_, Fig. 4(b,c), S3. We observe a transition from exponential to power-law behavior when *τ*_*s*_ becomes comparable to *T*_exp_ for a fixed *T*_*s*_. For small *τ*_*s*_, the potential landscape relaxes much faster than the time it takes to escape a potential well, resulting in exponential behavior as if the potential *U* (*x, s*) was static with *s* = *μ*_*s*_. For intermediate *τ*_*s*_, the first passage time distribution behaves as a truncated power law, with an exponential cut-off emerging for *τ*_*s*_ *< t < T*_exp_, Fig. S3. For *τ*_*s*_ sufficiently large, *τ*_*s*_ ∼ *T*_exp_, our adiabatic approximation holds and the simulations exhibit a power law tail. In Fig. 4(c), we keep *τ*_*s*_ large and vary *T*_*s*_ in the range [*T*_*x*_*/*4, 2*T*_*x*_]. The direct numerical simulations quantitatively recover the dependence of the power law exponent on the ratio between the fluctuations in *x* and *s*, Eq. 13, approaching *t*^−2^ as *T*_*s*_ → ∞.

### C. Long-range correlations and their finite-size corrections in slowly-driven metastable dynamics

In the previous section, we have shown that slowly varying barrier heights can give rise to power-law tails in the distribution of first passage times. Here, we extend these results to show that the correlation function also exhibits heavy tails, and that finite-size corrections give rise to the long-range anti-correlations observed in the worm behavior, Fig. 2(b-right). The (connected) normalized autocorrelation function of *x* is given by

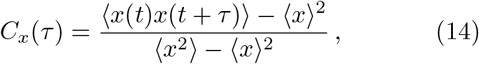

where ⟨· ⟩ represents the ensemble average over the invariant density. In our generic model of behavior, Eqs. 3,4, the long-time behavior of the correlation function of *x* is dominated by the first non-trivial eigenvalue of the Fokker-Planck operator, Λ_1_(*s*), which is proportional to the slowest hopping rate Λ_1_(*s*) ∝ *ω*(*s*) [33, 54],

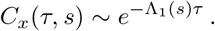

In the adiabatic limit, the correlation function of *x* can be obtained through a weighted average over the slowly fluctuating *ω*,

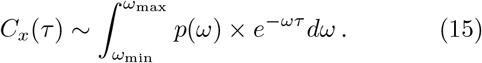

Notice that, compared to the expression for the first passage time distribution *f* (*t*), Eq. 6, the integrand is divided by *ω*^2^: one *ω* is dropped since *C*_*x*_(*t*) ≈*f* (*t, ω*)*/ω* and the other *ω* factor attributed to the effective number of observed transitions *ωT*_exp_ is also dropped since the correlation function is not simply determined by the transition events. Following the same steps as for *f* (*t*) (see Appendix B), we expect that

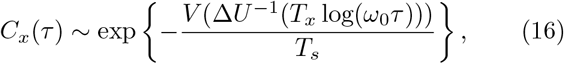

to the dominant order in the asymptotic approximation for large time delays *τ*. As for the FPTD, when *V* (*s*) and ∆*U* (*s*) are asymptotically equivalent, *C*_*x*_(*τ*) behaves as a power law 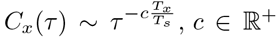. In this case, both the first passage time distribution *f* (*t*) ∼ *t*^*β*^ and the correlation function *C*_*x*_(*τ*) ∼ *τ*^*γ*^ have power-law tails, with exponents that are related by *γ* = *β* + 2.

To illustrate these results, we return to the example of the slowly forced overdamped dynamics on a double-well potential, Eq.12. The ergodic expectation would be that the correlation function should asymptote to 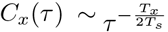. In particular, in the ergodic limit, we have ⟨*x*⟩ = 0 (since the potential is symmetric around *x*_*f*_ = 0) and the connected correlation function is simply given by its non-connected component 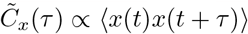. Indeed, if we measure the non-connected correlation function from numerical simulations of Eq. 12 (without subtracting the mean and normalizing by the variance, see Appendix A), we recover the theoretical expectation of power-law correlations for large *τ*_*s*_, Fig. 5(a).

**FIG. 5.**
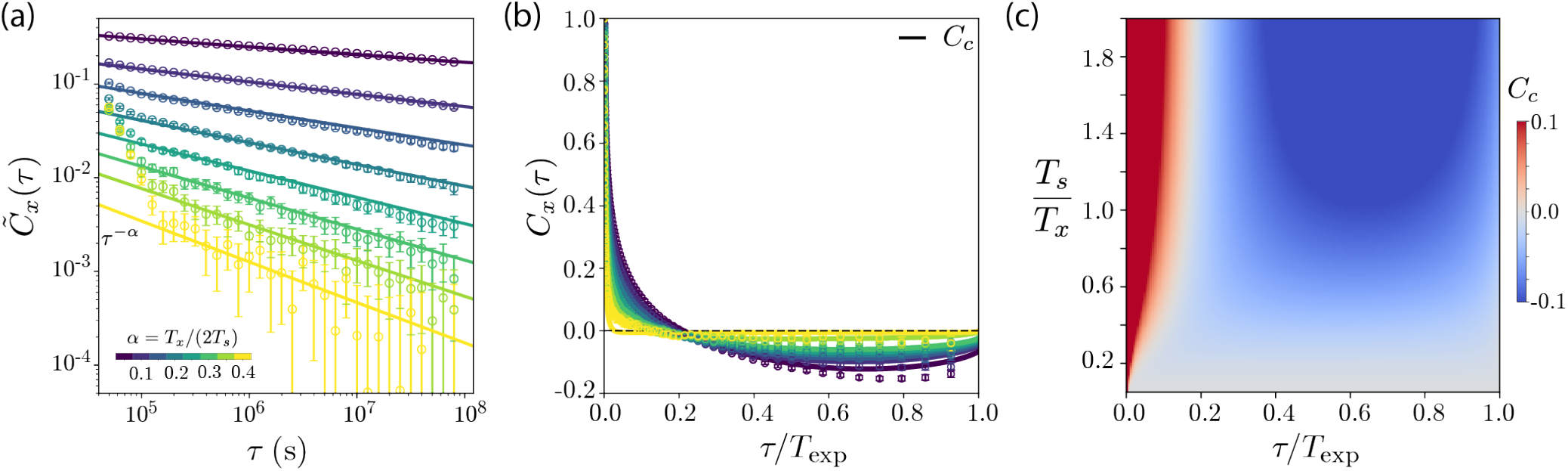
Power-law correlations and finite-size corrections in a slowly-driven double well potential. (a) Estimated non-connected correlation function 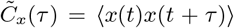 (see Appendix A) for the position of a particle in a double well potential driven on a timescale *τ*_s_ = 10^2^*T*_exp_. As predicted, the correlation function exhibits power-law tails 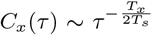 (solid lines). Error bars represent 95% confidence intervals across 50,000 simulations. Large *τ* estimates become increasingly challenging with the growth of the exponent. (b) Connected autocorrelation function, *C*_*x*_(*τ*), directly estimated through time averages (see Appendix A), for the position of a particle in a double well potential driven on a timescale *τ*_*s*_ = 10^2^*T*_exp_. Due to the existence of timescales comparable to the observation time *T*_exp_, the connected correlation function exhibits finite-size effects that drive the appearance of long-range anti-correlations, as predicted from the finite-size correction to the correlation function *C*_*c*_ [55] derived in Appendix D (solid lines). For both the empirical *C*_*x*_(*τ*) and *C*_*c*_ we normalize the correlation functions by dividing by their value at *τ* = 1 lag = 5 *×* 10^−4^*T*_exp_. Error bars represent 95% confidence intervals across 50,000 simulations. Finite-size correction to the correlation function as a function of *T*_*s*_ (see Appendix D). As we increase the temperature, the range of observed *ω* grows, and so do the deviations from the ergodic expectation, resulting in more apparent finite-size effects with clear anti-correlations (blue) appearing for large *T*_*s*_*/T*_*x*_. Conversely, for very small *T*_*s*_ the finite size effects become negligible due to the fact that the longest sampled *ω*^−1^ is much shorter than the experimental time scale *T*_exp_.

Notice, however, that in the empirical estimation of the connected correlation function, the expectation values are typically estimated by averaging in time (see Appendix A). In the presence of slow time scales, we expect that such temporal averages 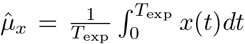 will deviate significantly from the ensemble average of *x* with respect to the invariant measure *π*(*x*), *μ*_*x*_ = *xπ*(*x*)*dx*. Such weak ergodicity breaking results in finite-size effects in the estimation of the connected correlation function from time series data [56]. In particular, when *x* relaxes to its steady state expectation value *μ*_*x*_ on time scales comparable to the observation time *T*_exp_, we expect that on average 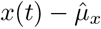will change signs as time progresses. This transient behavior results in apparent long-range anti-correlations, since on average 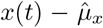 and 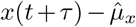 will have different signs for large *τ* [57]. Therefore, we expect that our derivation of Eq. 16 will deviate from the direct empirical estimation of the connected autocorrelation function, particularly when slow time scales are present in the dynamics. Indeed, when we estimate the connected correlation function directly through temporal averages (see Appendix A), we observe the appearance of long-range anti-correlations, Fig. 5(b). Importantly, using our analytical derivation of the non-connected correlation 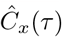 and the results of [55], we can derive an expression for the finite-size correction *C*_*c*_(*τ*) to the connected correlation (see Appendix D), that correctly approximates the behavior of the empirical estimate of *C*_*x*_(*τ*), Fig. 5(b).

As *T*_*s*_ becomes smaller, the finite-size effects become less apparent, reflecting the fact that the slowest generated timescales are closer to the mean hopping rate, Fig. 5(c). Conversely, finite-size effects become clearer the larger *T*_*s*_ is. In addition, for sufficiently small *τ*_*s*_ we observe that the non-connected correlation function exhibits exponential tails, which become a power law only when *τ*_*s*_ ∼ *T*_exp_, Fig. S4(a-left), which is the regime in which our adiabatic approximation holds. However, even when the tail of the correlation function is exponential, finite-size corrections are still apparent, as long as the exponential timescales are sufficiently long to be comparable to the measurement time scale, Fig. S4(a-right,b).

We therefore uncover that slowly-fluctuating energy landscapes generally give rise to long-range temporal correlations, which become anti-correlations due to finite-size effects. In addition, we find that even when the modulation of the potential landscape is fast, finite-size effects might still be apparent as long as the variation in the barrier height is sufficient to give rise to hopping rates that are comparable to the measurement time. Our theoretical results thus qualitatively recover the non-trivial correlations observed in foraging worms, Fig. 2(b-right).

## IV. DISCUSSION

We combine theory, numerics, and data analysis to show that the multiplicity of timescales inherent to animal behavior is sufficient to give rise to heavy-tailed first passage times and long-range correlations.

We start by analyzing the movement dynamics of the *C. elegans* nematode, where we build an effective reduced order model from high-resolution measurements of the animal pose, bridging from ∼ 0.1 s chaotic posture dynamics to ∼ 10 s stochastic hopping among “runs-and-pirouettes” [18]. The spatio-temporal dynamics of the animal’s pose results from a nontrivial combination of neural and biomechanical influences, for which building a first-principles microscopic model is extremely challenging. Nonetheless, we here obtain a one-dimensional stochastic differential equation that self-consistently captures the long-lived properties of the inferred dynamics, Fig. 1(d).

Technically speaking, we build an overdamped description of a partially-observed system with metastable dynamics, combining ideas from reduced order modeling [25, 58–60] and stochastic model inference [32, 61, 62]. Our contribution here is mostly conceptual: instead of assuming structure *a priori*, we reveal it by carefully analyzing the time series. We first found that time delays are required to recover minimal-memory evolution [9, 11], then leveraged transfer operators to discover that the system exhibits timescale-separated long-lived modes [11], and finally recovered a self-consistent effective overdamped Langevin description of the slow mode dynamics. This combination of first principles approaches, which is agnostic to the properties of the system under study, is what allowed us to build such a complete picture of a system for which physical intuition is elusive. We note, however, that each step in our framework could be enhanced by incorporating modern tools from machine learning (see, e.g., [63]), and we leave that for future work.

We find heavy-tailed first passage time distributions and correlation functions in the worm data, Fig. 1(d), that could not be captured by a model featuring autonomous dynamics only. This is perhaps inevitable: behavior is influenced by a wide range of biological mechanisms acting across a multitude of scales, and the fact that an autonomous Markovian description of the posture dynamics is predictive at all is surprising [64, 65]. In fact, while our framework tries to capture the effect of *hidden* variables through a delay embedding, slow non-ergodic modes would require a prohibitively large number of time delays to be properly encoded in the reconstructed state-space. To capture such non-ergodic modulations, we leverage our stochastic model as a locally-adiabatic approximation of the dynamics, and allow the parameters that define the effective potential landscape to change in time. This approach allows us to improve upon the predictions of the static model, and effectively capture the heavy-tailed first passage time statistics and the long-range correlations exhibited by the worm, Fig. 2.

In addition, we find that the slow modulation of the potential landscape effectively encodes the adaptation of the worm’s foraging strategy, biasing their random walk to search for food further away. In this way, we go beyond classical approaches to reduced-order modeling, recognizing the existence of non-ergodic fluctuations and introducing an explicitly non-stationary model to encode them.

Our results indicate that the heavy tails observed for *C. elegans* foraging behavior results from the slow adaptation of their search strategy. To test this experimentally, one could perturb the neural circuits responsible for the adaptation of “pirouette” rates, such as dopaminergic and glutamatergic signaling [34]. In the absence of adaptation, we expect that, while shorter timescale movements remain unaffected, the dwell times in the “run” and “pirouette” states would become exponentially distributed, rather than heavy-tailed, and that the correlation function would simply decay exponentially to zero on faster timescales.

The analysis of *C. elegans* data also suggests a general mechanism for the emergence of heavy-tailed first passage time distributions and long-range correlations in animal behavior. We investigate this by introducing a generic model in which the posture dynamics evolves on slowly fluctuating potential landscapes. We find that when non-ergodic fluctuations are sufficiently slow and strong, the first passage time distribution asymptotes to a power law with an exponent *f* (*t*) ∼ *t*^−2^, and otherwise exhibits corrections that depend on the ratio between the ergodic and non-ergodic fluctuations and the details of the dynamics. In addition, we find that estimates of the connected correlation function exhibit long-range anti-correlations due to finite size effects and that, in the absence of such effects, the correlations would be long-ranged, appearing scale-free when *V* (*s*) and ∆*U* (*s*) are asymptotically equivalent.

In the context of animal behavior, heavy-tailed distributions of first passage times (or run lengths, at constant speed) with an exponent *f* (*t*) ≈*t*^−2^ have been found extensively across multiple species, from bacteria [66], termites [67] and rats [68] to marine animals [69, 70], humans [71] and even fossil records [72]. In the context of search behavior, such observations have led researchers to hypothesize that Lévy-flights (with an exponent of -2) result in efficient search strategies and are thus evolutionarily favorable [73–77], although this view has been met with some controversy [78, 79]. Indeed, we here show that fat tails appear simply from the fact that the animal continuously adapts its behavior over time, leading to a broad distribution of “run” times (and thus run lengths). Therefore, such emergent behavior need not be fine-tuned by evolution *per se*, although one might argue that it is a by-product of the evolutionarily favorable ability to perform adaptive behavior.

Interestingly, the notion that time-dependent energy barriers can give rise to power-law waiting time distributions has also been used to explain observations in bacterial chemotaxis [80]. However, the analysis of [80] concerns only a particular limit of our derivation, in which *T*_*s*_→ ∞ and the distribution of hopping rates becomes uniform. Indeed, our analysis is more general: we consider the full dynamics of coupled overdamped Langevin equations, and predict corrections to the power-law behavior that go beyond the limits deployed in [80]. Another mechanism that has been proposed to explain the emergence of Lévy-flights in animal behavior is the existence of multiplicative noise terms in the dynamics [81, 82], and this notion has recently been used to explain the emergence of Lévy-flights in the collective behavior of midge swarms [83]. Our analysis is somewhat analogous to this argument. Indeed, Eqs. 3,4 give rise to an effectively colored multiplicative noise term for the quasistationary behavioral dynamics. However, our analysis also goes beyond that of [83], as we explicitly explore the dependency of the heavy tails on the relationship between the correlation time of the colored noise *τ*_*s*_ and the measurement time *T*_exp_, and between the additive and multiplicative noise terms.

Our starting point is an effective description of the long-time scale dynamics, Eqs. 3,4, and further work will be required to fully bridge between the microscopic dynamics and the emergent long time behavior, especially when a time scale separation is not evident, or when the adiabatic approximation does not hold. For example, we find that for intermediate values of 1 ≪ *τ*_*s*_ ≪*T*_exp_ and finite *T*_*s*_ the numerically estimated FPTD behaves as a truncated power law with an effective exponent that can be smaller than − 2. In this regime, the barrier heights fluctuate significantly before the particle hops. Intuitively, we expect that if barrier-crossing events become uncorrelated, the extra *ω* correction in the FPTD, coming from the increased probability of observing hopping events for larger *ω*, drops out, resulting in a FPTD with a dominant power law with an exponent of *t*^−1^, instead of *t*^−2^ [84]. In the opposite regime, we note that when *τ*_*s*_ *T*_exp_, it is the distribution of initial conditions (which we here assumed to be Boltzmann distributed) that determines the emergent behavior. This assumption holds if we consider that behavioral “individuality” is equivalent to having an extremely slow mode driving the dynamics *τ*_*s*_ ≫ *T*_exp_. This would mean that in finite observations from a population of conspecifics, different animals will exhibit a degree of “individuality” that matches the steady-state distribution of such long-lived modes. Indeed, such a relationship between interindividual variability and long-lived temporal variability in behavior has been observed in flies [85]. In this sense, when *τ*_*s*_ ≫*T*_exp_, our results are equivalent to explaining the emergence of heavy tails through inter-individual variability [86]. If such variability differs from the Boltzmann assumption, the heavy tails need to be corrected accordingly, following the steps of our derivation but with a corrected *p*(*s*).

In addition to heavy-tailed FPTDs, we also derived that when the barrier height is slowly-driven long-range correlations emerge, which become anti-correlations when the ergodic assumptions break and the temporal averages become plagued by finite-size effects. Notably, a recent arXiv preprint that followed the original arXiv version of this manuscript, has shown evidence for powerlaw correlations in the behavior of fruit flies [87]. These observations fit our theoretical predictions, and we argue that they might stem from non-ergodic internal states. Indeed, we expect there to be slow modes that evolve on timescales comparable to the 1 hour recordings used in [87], see [88, 89].

Power laws have been observed in a wide variety of systems, from solar flares [90, 91] to the brain [92] and different hypotheses have been put forward to explain their emergence (for a review see e.g. [93]). In disordered systems for example [94, 95], averaging over an exponential distribution of barrier heights can give rise to a broad distribution of waiting times. Note however, that while this mechanism is at its core analogous to the one presented here, ours relies on the temporal (rather than spatial) variation of barrier heights, resulting in distinct emergent behavior that depends directly on the measurement time scale *T*_exp_ (that sets the lowest hopping rate *ω*_min_) and the magnitude of the non-ergodic fluctuations.

With respect to power laws in biological systems and, in particular, in neuroscience, work inspired by phase transitions in statistical mechanics associates such power laws to “criticality” [96, 97], since models inferred from data appear to require fine-tuning of the parameters to a special regime between two qualitatively different “phases” (see, e.g., [98]). However, power laws can emerge without fine tuning and far from “criticality” [99], a clear example being Alder tails in hydrodynamics [100]. Here, we show how apparent “criticality” can emerge from the presence of slow non-ergodic drives, regardless of the details of the dynamics. Indeed, slow modes that evolve on time scales comparable to the observation time are challenging to infer from data, and can give rise to best-fit models that appear “critical”. While some of the arguments we have put forward have also been proposed to explain neural “criticality” [101–103], we here generalize to a wider range of model classes, using the framework of out-of-equilibrium statistical mechanics to explicitly connect the long time scale emergent behavior with the underlying effective fluctuations. In addition, unlike other approaches [102, 104], our framework does not require explicit external drives, but simply collective modes that evolve in a weakly non-ergodic fashion.

We have used a physics approach to shed light on biological phenomena, leveraging statistical mechanics as a framework for thinking about the effect of slowly-varying internal states on animal behavior. Simultaneously, the observations from animal behavior also inspired new physics, leading to general results regarding the emergence of heavy tails in slowly-driven potential landscapes, a result that we believe is relevant to a wide range of natural systems in chemistry, biology, or finance (see, e.g., [44–47, 50, 105, 106] and references therein).

## V. ACKNOWLEDGEMENTS

We thank Adrian van Kan, Stéphan Fauve, Federica Ferretti, Tosif Ahamed, Nicola Rigoli and Arghyadip Mukherjee for their comments. This work was partially supported by the LabEx ENS-ICFP: ANR-10-LABX-0010/ANR-10-IDEX-0001-02 PSL*. AC also acknowledges useful discussions at the Aspen Center for Physics, which is supported by National Science Foundation Grant PHY-1607611.

## APPENDIX A METHODS

### Software and data availability

Code for reproducing our results is publicly available: https://github.com/AntonioCCosta/fluctuating_potential. Data can be found in [107].

#### *C. elegans* foraging dataset

We used a previously-analyzed dataset [26], in which N2-strain *C. elegans* were tracked at *f* = 16 Hz [24]. Worms were grown at 20°*C* under standard conditions [108]. Before imaging, worms were removed from bacteria-strewn agar plates using a platinum worm pick, and rinsed from *E. coli* by letting them swim for 1 min in NGM buffer. They were then transferred to an assay plate (9 cm Petri dish) that contained a copper ring (5.1 cm inner diameter) pressed into the agar surface, preventing the worm from reaching the side of the plate. Recording started approximately 5 min after the transfer and lasted for 35 mins.

### Data-driven reduced order model of *C. elegans* foraging dynamics

Building upon previous work [9– 11], we extract a slow reaction coordinate that captures transitions between “runs” and “pirouettes” from the posture dynamics of *C. elegans*. The first step consists in performing a time-delay embedding of the instantaneous posture measurements to include short-term memory into an expanded maximally predictive state *X*_*K*∗_. The amount of time delays *K*^∗^ used to reconstruct the state space is chosen so as to maximize predictive information [10, 11]. In this way, all the dynamics that mix on a sufficiently fast timescale (compared to the measurement time) should be included in the state. We then partition the state space into a large number of discrete symbols through k-means clustering, and choose the number of partitions so as to preserve as much information as possible in the discretization [10, 11]. The outcome of the partitioning is a symbolic sequence, where each symbol *s*_*i*_ corresponds to a small region of the state space. We then build a Markov chain by counting transitions among state-space partitions separated by a timescale *τ, P*_*ij*_(*τ*) = *P* (*s*_*j*_(*t* + *τ*) *s*_*i*_(*t*)) ≈*e*^*ℒτ*^ [10, 11], effectively approximating the action of the Perron-Frobenius operator (see, e.g., [109]). The transition time *τ*^∗^ = 0.75 s was chosen so as to self-consistently capture the long-lived dynamics [10, 11]. The eigenfunctions of the Perron-Frobenius operator, and its adjoint, the Koopman operator, capture global patterns of the dynamics that relax to the steady-state distribution on different timescales. In particular, the slowest eigenfunctions of these operators offer optimal reaction coordinates that capture the slow dynamics of the system [59, 60, 110]. The slowest left eigenvector of the reversibilized transition matrix 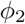 captures transitions among “runs” and “pirouettes” that *C. elegans* uses to forage [11]. We find the transition point 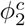 between “runs” and “pirouettes” by maximizing the overall coherence of the metastable states [10, 11], and recenter and rescale *ϕ*_2_ to have 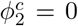 at the transition point and to have equally-spaced values within 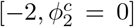 and 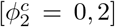. Finally, each symbol *s*_*i*_ assumes a particular value of *ϕ*_2_, and so we can translate the symbolic sequence into a stochastic time series *ϕ*_2_(*t*) that captures transitions between “runs” and “pirouettes”, see Fig. 1(a-c).

### Two-dimensional UMAP embedding of the reconstructed state space

We use the UMAP embedding [111] as a tool to visualize the maximally predictive states *X*_*K*∗_ of *C. elegans* posture dynamics [11]. In a nutshell, the UMAP algorithm searches for a low-dimensional representation of the data that preserves its topological structure. We use a publicly available implementation of the algorithm https://github.com/lmcinnes/umap, using Chebyshev distances, <monospace>n neighbors</monospace>=50 nearest neighbors and min dist=0.05 as the minimum distance.

### Stochastic model inference

The Kramers-Moyal expansion transforms the master equation for the dynamics of Eq. 1 into a Fokker-Planck equation

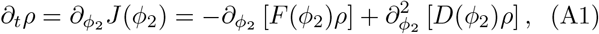

where *J* (*ϕ*_2_) is the current, and

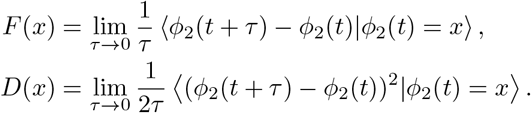

We use this expansion to estimate *F* (*ϕ*_2_) and *D*(*ϕ*_2_). In practice, given a time series *Y*_*t*_, we estimate the averages in the Kramers-Moyal expansion using a kernel approach [32],

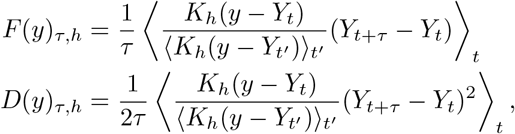

where *K*_*h*_(*z*) = *h*^−1^*κ*(*z/h*) and *κ* is the Epanechnikov kernel [112, 113],

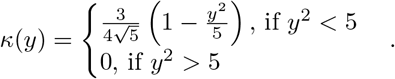

Importantly, the estimator has an explicit dependence on the time delay *τ* and the bandwidth *h*. First, as discussed in the main text, we choose *τ* long enough such that most of the temporal correlations in the noise have decayed to zero. It has been shown that *τ*^∗^ = 0.75 s gives an accurate first-order Markov model of the worm dynamics [11], and accordingly we find that a stochastic model inferred with *τ*^∗^ = 0.75 s yields nearly delta-correlated noise, Fig. S1(b). Given this time delay *τ*^∗^, we choose the bandwidth through the ∆-algorithm introduced in [32]. In essence, for each bandwidth *h* we estimate *F*_*τ*∗, *h*_ and *D*_*τ*∗, *h*_ and generate simulations with the estimated *F*_*τ*∗, *h*_ and *D*_*τ*∗, *h*_. From such simulations, we then re-infer the drift and diffusion from the simulated time series, obtaining 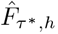 and 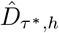. Finally, we compare the re-inferred drift and diffusion to the ones estimated directly from the time series,

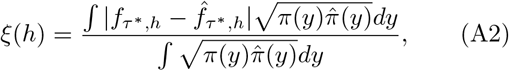

where *f* can be either *F* or *D, π*(*y*) is the steady-state distribution obtained from *F*_*τ*∗, *h*_ and *D*_*τ*∗, *h*_ and 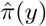 is the one obtained from 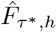 and 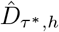. We choose *h*^∗^ as the first minimum of *ξ*(*h*) [32], locally minimizing the difference between original and reconstructed drift and diffusion coefficients and avoiding the trivial minimum that corresponds to *h*→ ∞ (which yields constant *F* and *D*). In Fig. S1(a) we plot the change in *ξ*(*h*) as a function of *h*, which reaches zero at around *h*^∗^ ≈0.1. We choose *h*^∗^ = 0.08 to infer the time series of *ϕ*_2_(*t*).

#### Non-stationary stochastic model inference

We proceed as before, but now infer 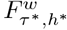 and 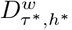 overlapping 5 min windows. The window length was chosen long enough to allow for equilibration of the “run” and “pirouette” dynamics, which has a mixing time of ≈ 3 s [11], but also short enough such that the steady-state distribution remains approximately constant.

#### Reconstructing an effective potential landscape

From the Fokker-Planck equation, Eq. A1, with natural boundary conditions, *J* (*ϕ*_2_) = 0, we can obtain the steady-state solution *ρ* = *π*, satisfying ∂_*t*_*π* = 0, as,

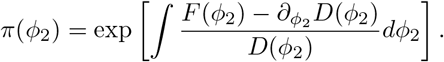

Writing the steady-state distribution as a Boltzmann factor [114],

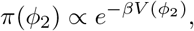

with *β* = 1, we can identify an effective potential land-scape *V* (*ϕ*_2_),

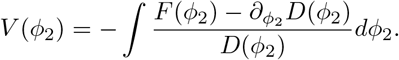

The same approach applies to the time-dependent stochastic model, where each window has its own local steady-state, and the effective potential landscape is time dependent,

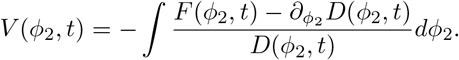

#### Stochastic model simulations of *ϕ*_2_

We simulate the dynamics using an Euler scheme with the same sampling time as the data *δt* = 1*/*16 s. For the non-autonomous model, we take 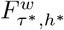 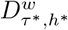 of the window with a center closest to the sampled time point.

#### Fine-scale Markov model simulations

As in [11], we simulate symbolic sequences by sampling the next state according to the condition probability distribution *P* (*s*_*j*_(*t*+*τ*^∗^) | *s*_*i*_(*t*)), which is simply the *i*-th row of *P*_*ij*_(*τ*^∗^). From this symbolic sequence, we can then obtain a simulated time series of *ϕ*_2_(*t*) sampled on a timescale *τ*^∗^.

#### Estimating the first passage time distributions in *C. elegans* foraging dynamics

We estimate the time spent either performing a “run” or a “pirouette” by identifying segments where *ϕ*_2_(*t*) *<* 0 (runs) or *ϕ*_2_(*t*) *>* 0 (pirouettes). To remove short-time fluctuations we sub-sample the data and the simulated time series by *τ*^∗^*/*2.

#### Empirical estimate of the connected auto-correlation function

We estimate the connected autocorrelation function from *M* time traces at each lag *τ* = *lδt*, as

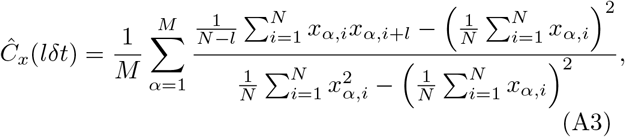

where *x*_*α,i*_ is the *i*-th frame of the *α* trace with length *N* = *T*_exp_*/δt*.

#### Estimating the first passage time distribution of a Poisson process with varying hopping rates

We sample *s* according to the Boltzmann distribution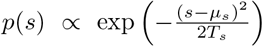, and convert it to a hopping rate *ω*(*s*) by numerically integrating the backward Kolmogorov equation, Eq. 10. We then sample first passage time events according to Eq. 5, until reaching the measurement timescale *T*_exp_. We repeat this process 50,000 times, and collect the statistics of waiting times to build a normalized histogram of first passage times with logarithmic bins, which we show in Fig. 3.

#### Estimating the first passage time distribution in the slowly-driven double well potential

We generate 10,000 simulations of Langevin dynamics of Eq. Eq. 12, through an Euler-scheme with a sampling time of *δt* = 10^−3^ s for *T*_exp_ = 10^7^ s. We then vary *τ*_*s*_ in the range [10^−4^*T*_exp_, 10^4^*T*_exp_] and *T*_*s*_ in the range [*T*_*x*_*/*4, 2*T*_*x*_], where *T*_*x*_ = 10^−3^ and 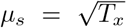. The initial conditions *x*(0) are sampled randomly either as *x*(0) = 1 and *x*(0) = −1 with equal probability and 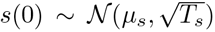 is sampled according to the Boltzmann distribution. From the simulations of *x*(*t*), we then estimate the first passage time distribution by first identifying all the segments, [*t*_0_, *t*_*f*_], in which *t*_0_ corresponds to the first time *x* returns to *x*_0_ = *±* 1 after reaching *x*_*f*_ = 0, and *t*_*f*_ is the time first to reach *x*_*f*_ = 0 after *t*_0_. Finally, we build a normalized histogram of first passage times with logarithmic bins, which we show in Figs. 4,S3.

#### Estimating the autocorrelation functions in the slowly-driven double well potential

We generate 50,000 simulations of Langevin dynamics of Eq. 12, through an Euler-scheme with an initial sampling time of *dt* = 0.2 s that is downsampled to *δt* = 100 s for *T*_exp_ = 10^8^ s, with *T*_*x*_ = 10^−2^, 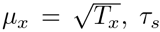, *τ*_*s*_ sampled in the range [10^−2^*T*_exp_, 10^2^*T*_exp_] and *T*_*s*_ sampled in the range [1.25*T*_*x*_, 10*T*_*x*_]. We then estimate the connected autocorrelation function from the simulations using Eq. A3. The non-connected correlation function is estimated as,

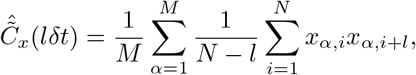

and then normalized by dividing by 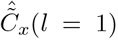. The finite-size corrections to the correlation function are detailed in Appendix D.

## APPENDIX B FIRST PASSAGE TIME DISTRIBUTION IN SLOWLY FLUCTUATING POTENTIAL LANDSCAPES

We here derive the expression for the first passage time distribution (FPTD) in a fluctuating potential landscape. As discussed in the main text, we consider the adiabatic limit in which the FPTD can be approximated by

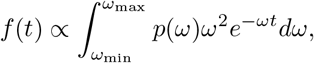

where *ω* = *ω*_0_ exp ∆*U* (*s*)*/T*_*x*_ *s*(*ω*) = ∆*U*^−1^(− *T*_*x*_ log (*ω/ω*_0_)), and *ω*_0_ is a typical (fast) frequency of the hopping dynamics [115]. The distribution *p*(*ω*) obeys *p*(*ω*)*dω* = *p*(*s*)*ds*, where *p*(*s*) ∝ exp *{*−*V* (*s*)*/T*_*s*_*}*, and is thus given by

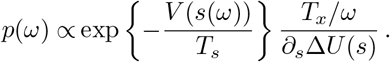

Plugging this into Eq. 6, we get

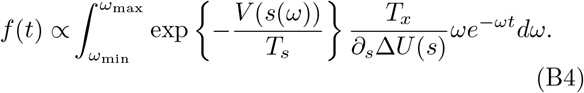

The exponential factor *e*^−*ωt*^ restricts the contributions to *ω* ∼ 1*/t*, which motivates the change of variable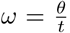. The above integral is then recast in the form

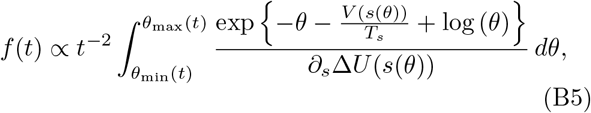

where 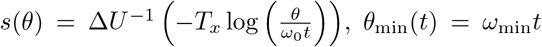 and *θ*_max_(*t*) = *ω*_max_*t*.

To grasp the structure of the integral, it is convenient to consider first the special case where *V* and ∆*U* can be written as a power series expansion *V* (*s*) ∼ *as*^*n*^ and ∆*U* (*s*) ∼ *bs*^*n*^, *a, b* ∈ℝ with an equal dominant (at large values of the argument, see below) exponent *n*. The integral reduces then to the form

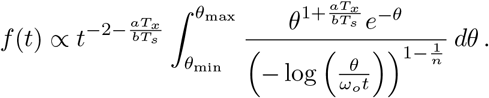

It remains to verify that the time dependencies at the denominator of the integrand and the limits of integration do not spoil the behavior at large times. This is verified by noting that the numerator of the integrand has the structure of an Euler-Γ function of order 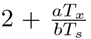. The numerator of the integrand has its maximum at 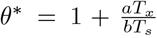, decays over a range of values of order unity (we consider^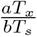^ to be small or of order unity) and vanishes at the origin. In that range, the argument of the power at the denominator log (*ω*_0_*t*) − log (*θ*) log (*ω*_0_*t*), which yields the final scaling with subdominant logarithmic corrections

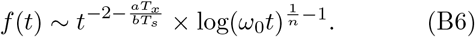

To complete the argument, we note that the time dependency of *θ*_min_ is not an issue as long as values *θ ~ O*(1) are in the integration range. In practice, this means that the minimum hopping rate *ω*_min_ should be comparable to (or larger than) the measurement time, 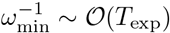.

Before moving to the general case, two remarks are in order. First, for *ω*_0_*t* ≫ 1 the functions *V* and ∂_*s*_∆≫ *U* that appear in Eq. B5 have their argument *s* ≫ 1. The dominant behavior of the two functions should then be understood for large values of their arguments. Second, the denominator ∂_*s*_∆*U* could *a priori* be included in the exponential at the numerator but this does not modify our conclusion. It is indeed easy to verify that the maximum *θ*^∗^ and the decay range would not be shifted at the dominant order (and this holds also for the general case considered hereafter).

We can now consider the general case with different dominant exponents *V* (*s*) ∼ *as*^*n*^ and ∆*U* (*s*) ∼ *bs*^*k*^, *a, b*∈ ℝ. The argument of the exponential in Eq. B5

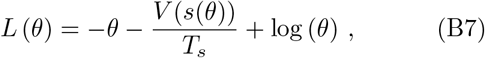

has its maximum at *θ*^∗^, defined by the implicit equation

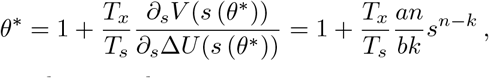

where we have used

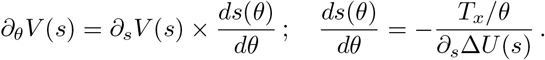

For *n < k*, the maximum *θ*^∗^ *≃*1 (as *s* 1) and the integrand decays in a range of order unity. Indeed, the dominant order of the derivatives ∂^*p*^*L* (*p*≥ 2) at *θ* = *θ*^∗^ coincide with those of log (*θ*). It follows that *L*(*θ*) − *L*(*θ*^∗^) ≃log (*θ/θ*^∗^) − (*θ* − *θ*^∗^). The resulting integral over *θ* is an Euler Γ-function of order two, which indeed forms at values *O*(1). In that range, 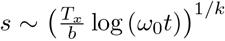 and the integral is then approximated by exp {*L* (*θ*^∗^)}, so *f* (*t*) becomes

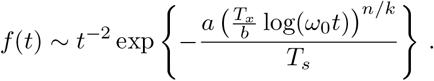

The factor at the denominator in Eq. B5 is 𝒪 [exp (1*/k* 1) log[log(*ω*_0_*t*)]] and thus of the same order as terms that we have discarded in our approximation so we neglect it as well. Since the integral over *θ* forms for values *O*(1), the constraint on the minimum hopping rate is the same as for the *n* = *k* case, i.e., 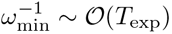.

For *n > k*, the maximum *θ*^∗^ ∼ (log *ω*_0_*t*)^*n/k*−1^, which is now large. The dominant order of the derivatives ∂^*p*^*L* (*p* ≥ 2) at *θ* = *θ*^∗^ is given by (− 1)^*p*−1^ (*p* − 1)! (*θ*^∗^)^−(*p*−1)^, that is they coincide with those of *θ*^∗^ log (*θ*). It follows that *L*(*θ*) − *L*(*θ*^∗^) ≃*θ*^∗^ [log (*θ/θ*^∗^) − (*θ* − *θ*^∗^) */θ*^∗^]. The resulting integral over *θ* is an Euler Γ-function of (large) argument *θ*^∗^ + 1 : its value is approximated by Stirling formula, which yields 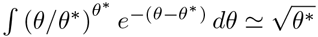. The 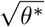 reflects the fact that the integral forms around the maximum at *θ*^∗^ of the integrand over a range 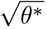, which implies that the approximation 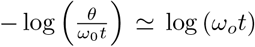 still holds, as in the previous cases *n* ≤ *k*. The 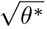, as well as the log (*ω*_0_*t*)^1*/k*−1^ coming from the denominator in Eq. B5, are subdominant with respect to terms that we have neglected in the expansion of *L*. We therefore discard them from our final approximation for *n > k* :

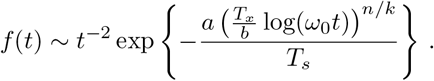

Since the integral over *θ* forms for values *𝒪(*(log *ω*_0_*t*)^*n/k*−1^) ≫ 1, the condition 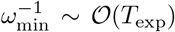 ensures *a fortiori* that the finite value of *ω*_min_ does not affect the above result.

Discarding subdominant terms, in all three cases we thus get the general expression we present in the main text,

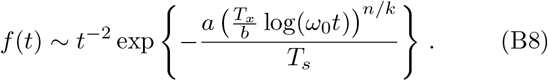

To verify the validity of the above arguments, we show in Fig.S5 how, to the dominant order, asymptotic predictions agree with a detailed numerical integration of Eq. B4 for ∆*U* (*s*) = *s*^*k*^ and *V* (*s*) = *s*^*n*^.

## APPENDIX C CORRELATION FUNCTIONS IN SLOWLY FLUCTUATING POTENTIAL LANDSCAPES

We here derive the expression for the correlation function in a fluctuating potential landscape. In general, the autocorrelation function can be expressed as a sum over exponential functions [33, 54],

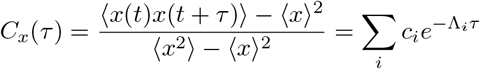

where Λ_*i*_ (Λ_1_ *<* Λ_2_ *<* …) are the eigenvalues of the Fokker-Planck operator and Σ_*i*_ *c*_*i*_ = 1. Since we are interested in the long-term behavior of systems with energy barriers that can fluctuate over time, we assume that the large *τ* behavior of the correlation function asymptotes to

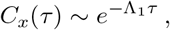

where Λ_1_ is the first non-trivial eigenvalue, which captures the longest-lived dynamics in the system. In addition, we assume that there is always a deeper well, with an escape rate *ω*, that dominates the long-lived dynamics. In this case, Λ_1_ ∝ *ω* and we have,

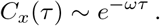

As previously discussed, we take the adiabatic approximation to derive the asymptotic behavior of the correlation function in the presence of slow non-ergodic modulation of the potential landscape. In particular, we obtain a weighted average of the correlation function over multiple realizations of *ω*(*s*), yielding

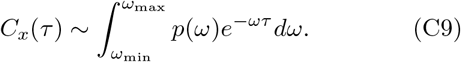

Note that in comparison with Eq. 6, besides dropping an *ω* factor due to the difference between *f* (*t, ω*), Eq. 5, and *C*_*x*_(*τ*), we also do not need to take into account the extra factor of *ω* coming from the finite observation time, which in the case of the estimation of first passage times biases the probability density in a manner that is proportional to *ω*. In the case of the correlation function, the dynamics of *x* is exposed to modulations in *s*, regardless of *ω*(*s*). Following the same steps as before, we consider that *V* and ∆*U* can be written as a series expansion with dominant terms *V* (*s*) ∼ *as*^*n*^ and ∆*U* (*s*) ∼ *bs*^*k*^, *a, b*∈ ℝ. In this case, we find that to dominant order

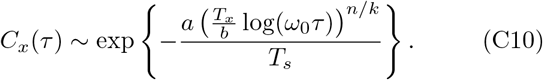

Notably, when *n* = *k*, we obtain power law correlations with an exponent that depends on the ratio of temperatures,

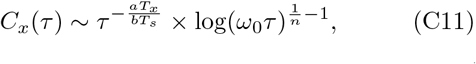

where we have included sub-dominant corrections. In particular, we find that when *n* = *k* both the first passage time distribution *f* (*t*) ∼ *t*^*β*^ and the correlation function *C*_*x*_(*τ*) ∼ *τ*^*γ*^ have power-law behavior at large times, and that the exponents are related by *γ* = *β* + 2. In addition, when *T*_*s*_ → ∞ correlations decay slowly as *C*_*x*_(*τ*) ∼ log(*ω*_0_*τ*)^1*/n*−1^.

## APPENDIX D FINITE-SIZE CORRECTION TO THE CORRELATION FUNCTION

When estimating the correlation function from a collection of *M* finite time traces sampled at *δt* and with length *N* = *T*_exp_*/δt*, we compute,

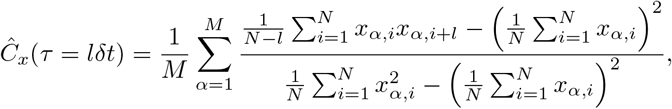

where *x*_*α,i*_ is the *i*-th frame of the *α* trace. Assuming that the finite-size corrections to the correlation function are dominated by corrections to the mean value (and not the variance), we can leverage the derivation of Desponds et al. [55] to obtain an expression for the finite-size corrections to the correlation function from the non-connected correlation function 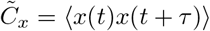,

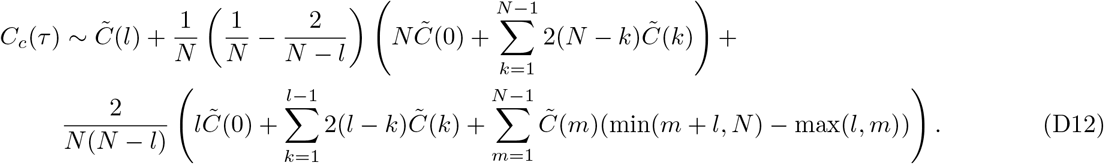

As detailed in the main text, as a case study we take the overdamped dynamics for the position *x* of a particle in a symmetric double well potential, for which the barrier height can fluctuate according to a slow parameter *s*, Eq. 12. The time scale separation between the hopping events and the relaxation to the well means that the correlation function is dominated by the first non-trivial eigenvalue, 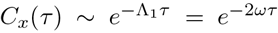, where we take Λ = 2*ω* due to the fact that the potential wells have the same depth [33]. Taking *V* (*s*) ∼ *s*^2^*/*2 and ∆*U* (*s*) = *s*^2^ we obtain that, in the asymptotic large *τ* limit, 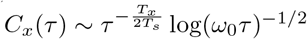, Fig. 5(a).

To obtain an accurate estimate of the correlation function for all *τ*, we go beyond the asymptotic approximation and numerically integrate

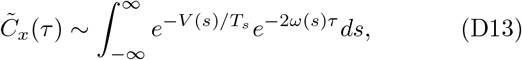

where *ω*(*s*) can be estimated directly by integrating the Kolmogorov backward equation. At large *τ*, the numerical integration of 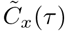 matches the asymptotic behavior 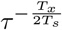. Plugging Eq. D13 into Eq. D12, we obtain the correction to the autocorrelation function *C*_*c*_(*τ*) presented in Figs. 5(b,c).

## SUPPLEMENTAL MATERIAL

**FIG. S1.**
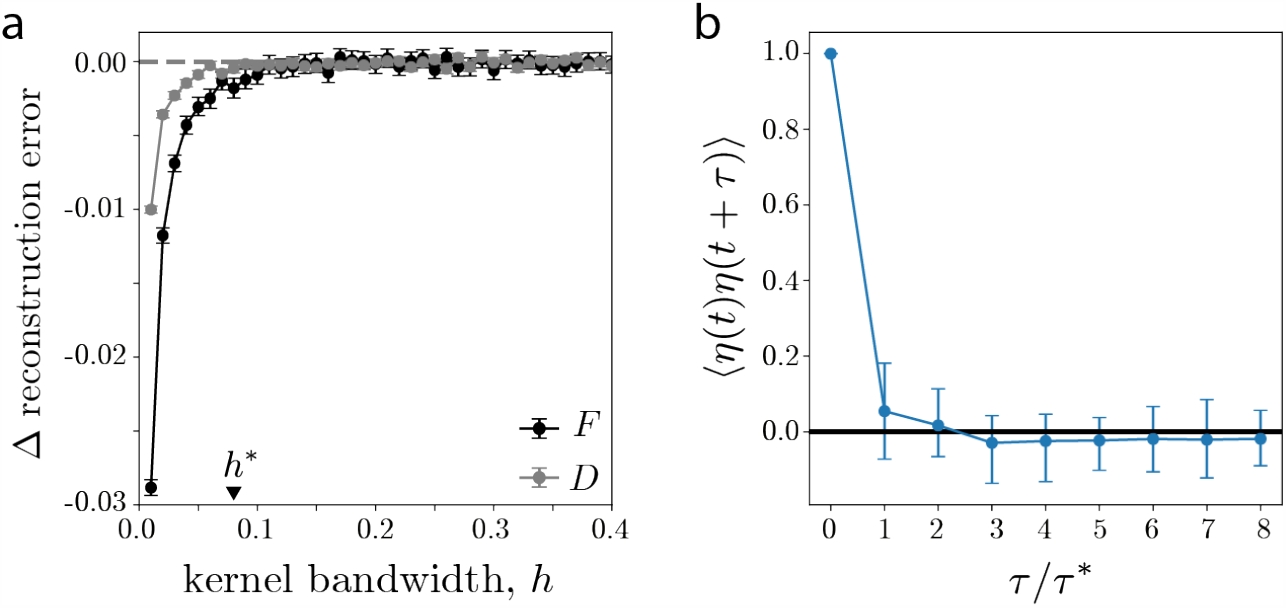
Details of the stochastic model inference. (a) Change in reconstruction error ∆*ξ* = *ξ*(*h* + *δh*) − *ξ*(*h*) of the drift *F* and diffusion *D* coefficients depending on the bandwidth of the kernel used to perform the Kramers-Moyal averages [32] (see Appendix A). For each value of bandwidth *h*, we simulate trajectories using the estimated drift and diffusion coefficients. We then re-infer the drift and diffusion coefficients from the simulated trajectories and compare them against the parameters obtained from the original time series to get the reconstruction error, Eq. A2 (∆-algorithm in [32]). We chose *h*^∗^ = 0.08 as the lowest *h* value when the reconstruction error stops changing (when ∆ reconstruction error ≈0). (b) Autocorrelation function of the residuals *η*(*t*) after fitting Eq. 1 to time series of *ϕ*_2_(*t*). At the sampling time *τ*^∗^ the noise decorrelates, thus justifying the white noise approximation. Error bars correspond to 95% confidence intervals bootstrapped over 1000 simulations of randomly sampled worms.

**FIG. S2.**
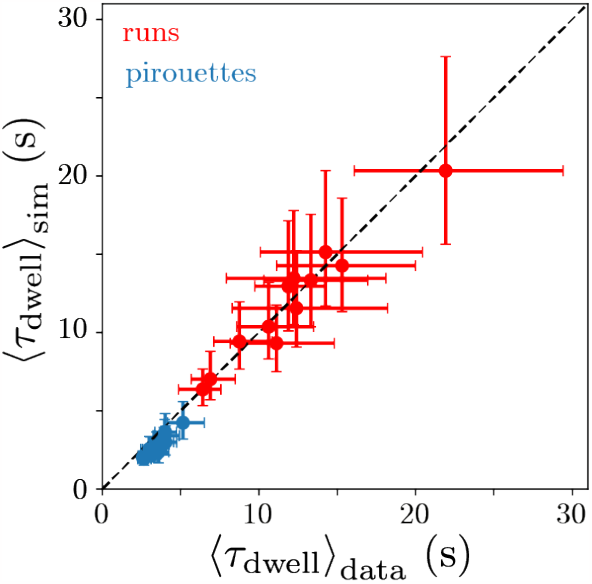
The simulations from the stochastic model accurately predict the average time spent in a given behavioral state. Each point corresponds to a worm in a particular state, and the error bars correspond to 95% confidence intervals bootstrapped over events.

**FIG. S3.**
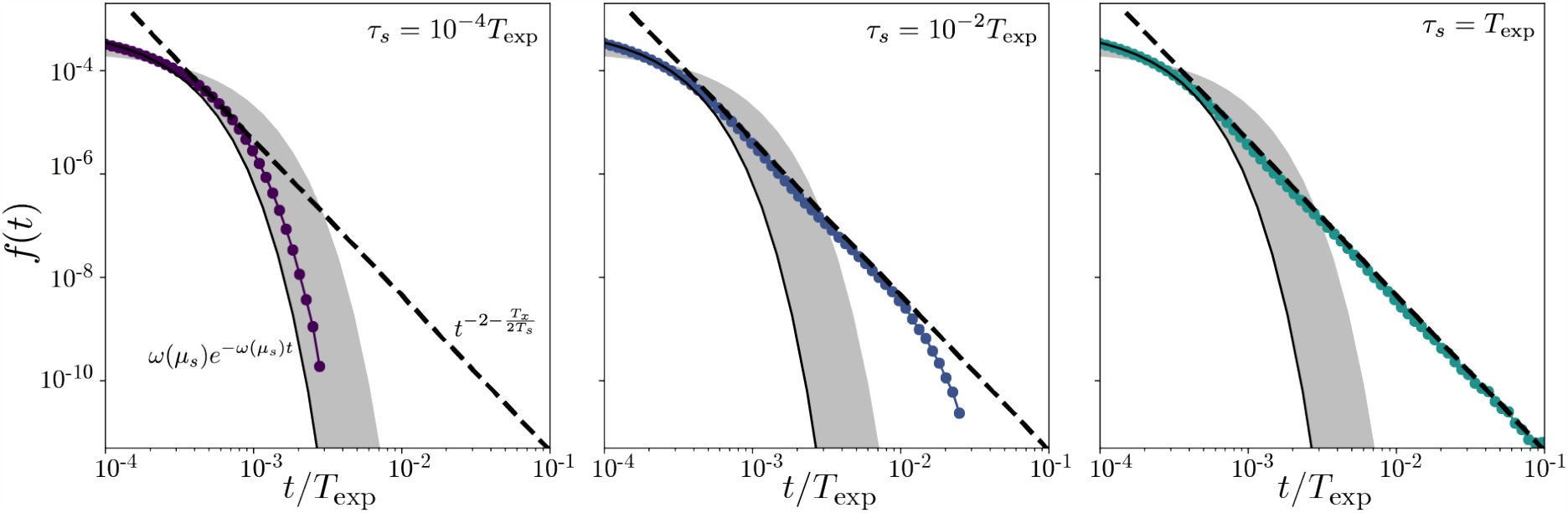
Emergence of power-law tails in the first passage time distributions for the slowly-driven double well dynamics of Eq. 12 as a function of *τ*_*s*_. For *τ*_*s*_ ≪ *T*_exp_ (left) the potential relaxes to its mean value faster than the hopping timescale, resulting in exponential behavior with a decay rate corresponding to the mean value *ω*(*μ*_*s*_). The black dashed line and gray shaded area correspond to *f* (*t, ω*) = *ωe*^−*ωt*^ with *ω* = *ω*(*μ*_*s*_) and 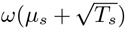 respectively. As *τ*_*s*_ increases, the regime in which we observe power law behavior grows, and for intermediate *τ*_*s*_ we obtain a truncated power law with an exponential tail starting at *t* ∼ *τ*_*s*_ (middle). Finally, when *τ*_*s*_ = *T*_exp_ the measured tail of the distribution is power-law distributed (right). The black dashed line corresponds to our prediction 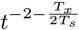 and *T* = *T /*2.

**FIG. S4.**
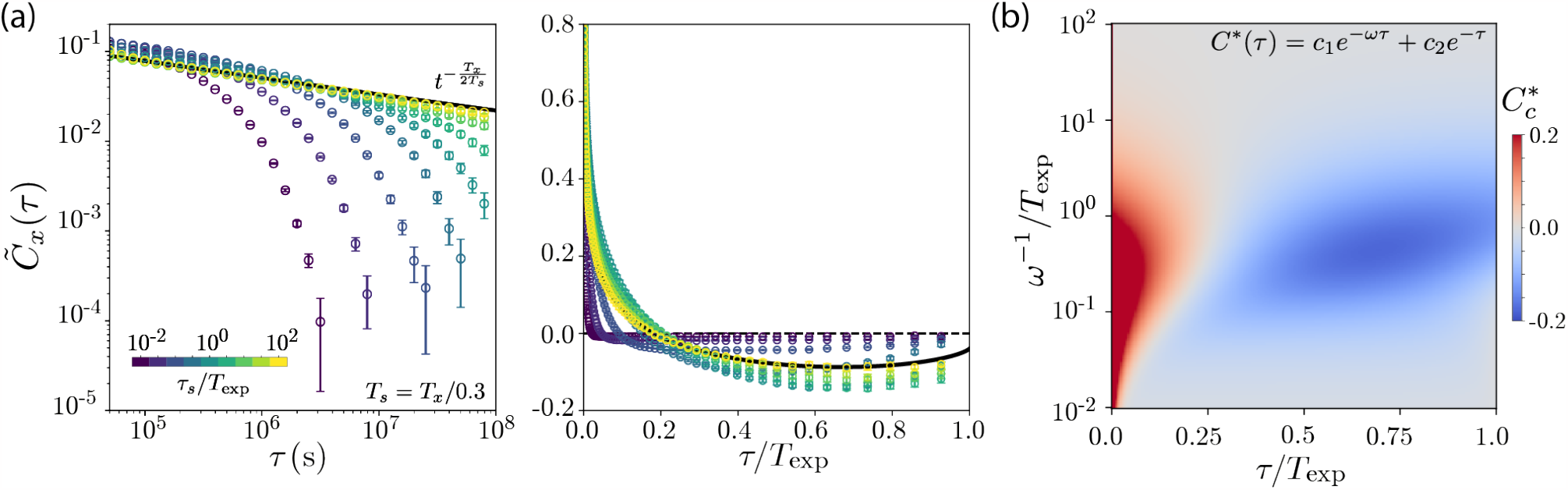
Dependence of the autocorrelation function on *τ*_*s*_. (a-left) Non-connected correlation function for *T*_*s*_ = *T*_*x*_*/*0.3 and varying *τ*_*s*_. When *τ*_*s*_ is small, the tail of the correlation function is exponential, with a decay rate that is given by the maximum barrier height attained at this temperature. As *τ*_*s*_ grows, the variation of the potential landscape becomes slower and slower and our asymptotic approximation correctly predicts the emergent power law behavior at large *τ*_*s*_. (a-right) Connected correlation function for *T*_*s*_ = *T*_*x*_*/*0.3 and varying *τ*_*s*_, estimated directly from the time series data. Our adiabatic approximation correctly predicts the correction to the correlation function for large *τ*_*s*_. We normalize both correlation functions by their value at *τ* = 1 lag = 5 *×* 10^−4^*T*_exp_. Notably, the correlation function exhibits finite-size corrections even when it has exponential tails, as long as the timescale of the exponential decay is comparable to the measurement time scale. Error bars represent 95% confidence intervals bootstrapped across 50,000 simulations. (b) We examine the finite-size correction to the correlation function when there is no time dependence to the hopping rate *ω*. For a system with a single fixed energy barrier, we would expect that the correlation function would be given by *C*_*x*_(*τ*) = *c*_1_*e*^−*ωτ*^ + *c*_2_*e*^−*τ*^ [33, 54], where Σ_i_*c*_*i*_ = 1 and we take the intrinsic time scale of relaxation to a well to unity without loss of generality. As expected, even when the correlation function has exponential tails we observe the appearance of finite-size effects when 0 ≪ *ω*^−1^ ;S ≲*T*_exp_. Notably, when *ω*^−1^→ ∞ these finite-size effects are less apparent since only the short timescale survives. In contrast, when we allow the hopping rate to fluctuate in time we effectively generate a continuum of time scales such that, even when *T*_*s*_→ ∞, finite-size effects are still apparent, Fig. 5(c).

**FIG. S5.**
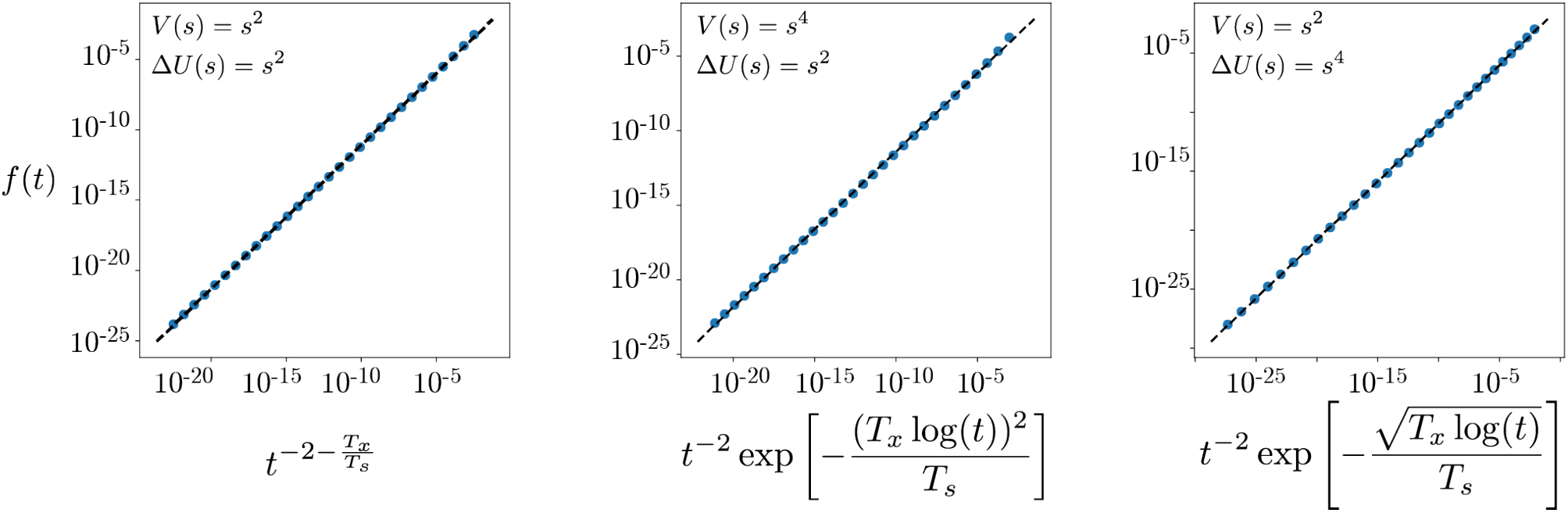
Numerical integration of *f* (*t*) for different choices of *V* (*s*) and ∆*U* (*s*), compared to the asymptotic approximation of Eq. B8 (black dashed line) with *T*_*x*_ = 0.1 and *T*_*s*_ = 0.2. We numerically integrate Eq. B4 with ∆*U* (*s*) = *s*^*k*^ and *V* (*s*) = *s*^*n*^, through a Riemann sum using the midpoint rule from *ω*_min_ = 5 *×* 10^−10^ to *ω*_max_ = 1 with ∆*ω* = 10^−9^.

## References

[1] T. D. Pereira, J. W. Shaevitz, and M. Murthy, Nature Neuroscience 23, 1537 (2020).

[2] M. W. Mathis and A. Mathis, Current Opinion in Neurobiology 60, 1 (2020).

[3] L. Hebert, T. Ahamed, A. C. Costa, L. O’Shaughnessy, and G. J. Stephens, PLoS Computational Biology 17, e1008914 (2021).

[4] G. J. Berman, BMC Biol. 16, 10.1186/s12915-018-0494-7 (2018).

[5] J. P. Sethna, Statistical Mechanics: Entropy, Order Parameters and Complexity, first edition ed. (Oxford University Press, Great Clarendon Street, Oxford OX2 6DP, 2006).

[6] J. Crank, The Mathematics of Diffusion, Oxford science publications (Clarendon Press, 1979).

[7] S. Brenner, Genetics 77, 71 (1974).

[8] C. I. Bargmann and E. Marder, Nat Methods 10, 10.1016/j.cub.2012.01.061 (2013).

[9] T. Ahamed, A. C. Costa, and G. J. Stephens, Nature Physics 17, 275 (2021).

[10] C. Costa, T. Ahamed, D. Jordan, and G. J. Stephens, Chaos: An Interdisciplinary Journal of Nonlinear Science 33, 023136 (2023).

[11] C. Costa, T. Ahamed, D. Jordan, and G. J. Stephens, A markovian dynamics for C. elegans behavior across scales (2023), arXiv:2310.12883 [physics.bio-ph].

[12] F. Takens, in Dynamical Systems and Turbulence, Warwick 1980, edited by D. Rand and L.-S. Young (Springer Berlin Heidelberg, Berlin, Heidelberg, 1981) pp. 366–381.

[13] G. Sugihara and R. M. May, Nature 344, 734 (1990).

[14] T. Sauer, J. A. Yorke, and M. Casdagli, Journal of Statistical Physics 65, 579 (1991).

[15] J. Stark, Journal of Nonlinear Science 9, 255 (1999).

[16] J. Stark, D. S. Broomhead, M. Davies, and J. Huke, Journal of Nonlinear Science 13, 519 (2003).

[17] J. T. Pierce-Shimomura, T. M. Morse, and S. R. Lockery, J. Neurosci. 19, 9557 (1999).

[18] M. Fujiwara, P. Sengupta, and S. L. McIntire, Neuron 36, 1091 (2002).

[19] H. C. Berg, E. coli in motion, Biological and medical physics series (Springer, New York, 2004).

[20] G. J. Berman, D. M. Choi, W. Bialek, and J. W. Shaevitz, J. Royal Soc. Interface 11, 1 (2014).

[21] B. Wiltschko, M. J. Johnson, G. Iurilli, R. E. Peterson, J. M. Katon, S. L. Pashkovski, V. E. Abraira, R. P. Adams, and S. R. Datta, Neuron 88, 1121 (2015).

[22] R. E. Johnson, S. Linderman, T. Panier, C. L. Wee, E. Song, K. J. Herrera, A. Miller, and F. Engert, Current Biology 30, 70 (2020).

[23] X. Brown and B. de Bivort, Nature Physics 14, 653 (2018).

[24] O. D. Broekmans, J. B. Rodgers, W. S. Ryu, and G. J. Stephens, eLife 5(e17227), 10.7554/eLife.17227 (2016).

[25] G. Froyland, G. A. Gottwald, and A. Hammerlindl, SIAM Journal on Applied Dynamical Systems 13, 1816 (2014).

[26] G. J. Stephens, B. Johnson-Kerner, W. Bialek, and W. S. Ryu, PLoS Comput. Biol. 4, e1000028 (2008).

[27] H. Mori, Progress of Theoretical Physics 33, 423 (1965).

[28] R. Zwanzig, Journal of Statistical Physics 9, 215 (1973).

[29] A. Rupe, V. V. Vesselinov, and J. P. Crutchfield, New Journal of Physics 24, 103033 (2022).

[30] This timescale is typically referred to as the Markov-Einstein scale [61, 116].

[31] We use the Itô interpretation of the stochastic dynamics (see, e.g., [117]).

[32] D. Lamouroux and K. Lehnertz, Physics Letters A 373, 3507 (2009).

[33] H. Risken and H. Haken, The Fokker-Planck Equation: Methods of Solution and Applications Second Edition (Springer, 1989).

[34] T. Hills, P. J. Brockie, and A. V. Maricq, Journal of Neuroscience 24, 1217 (2004).

[35] J. M. Gray, J. J. Hill, and C. I. Bargmann, Proceedings of the National Academy of Sciences 102, 10.1073/pnas.0409009101 (2005).

[36] L. C. Salvador, F. Bartumeus, S. A. Levin, and W. S. Ryu, Journal of the Royal Society Interface 11, 10.1098/rsif.2013.1092 (2014).

[37] J. Calhoun, A. Tong, N. Pokala, J. A. J. Fitzpatrick, T. O. Sharpee, and S. H. Chalasani, Neuron 86, 428 (2015).

[38] I. Hums, J. Riedl, F. Mende, S. Kato, H. S. Kaplan, R. Latham, M. Sonntag, L. Traunmüller, and M. Zimmer, eLife 5, 1 (2016).

[39] S. W. Flavell, D. M. Raizen, and Y.-J. You, Genetics 216, 315 (2020).

[40] I. Bargmann, BioEssays 34, 458 (2012).

[41] E. Marder, Neuron 76, 1 (2012).

[42] S. W. Flavell, N. Gogolla, M. Lovett-Barron, and M. Zelikowsky, Neuron 110, 2545 (2022).

[43] P. Hänggi, P. Talkner, and M. Borkovec, Rev. Mod. Phys. 62, 251 (1990).

[44] A. Szabo, K. Schulten, and Z. Schulten, The Journal of Chemical Physics 72, 4350 (1980).

[45] S. Condamin, V. Tejedor, R. Voituriez, O. Bénichou, and J. Klafter, Proceedings of the National Academy of Sciences 105, 10.1073/pnas.0712158105 (2008).

[46] O. Bénichou and R. Voituriez, Physics Reports 539, 225 (2014).

[47] R. Chicheportiche and J.-P. Bouchaud, Some applications of first-passage ideas to finance, in First-Passage Phenomena and Their Applications (World Scientific, 2014) Chap. 1, pp. 447–476.

[48] S. Grebenkov, Journal of Physics A: Mathematical and Theoretical 48, 013001 (2014).

[49] P. Hänggi, Chemical Physics 180, 157 (1994).

[50] O. Bénichou, T. Guérin, and R. Voituriez, Journal of Physics A: Mathematical and Theoretical 48, 163001 (2015).

[51] A. Godec and R. Metzler, Phys. Rev. X 6, 041037 (2016).

[52] N. G. Van Kampen, Stochastic processes in physics and chemistry (North-Holland, Amsterdam, 1981).

[53] We note that this is generally true even for fixed initial conditions as long as τs < Texp. If τs Texp, then p(ω) is primarily defined by the distribution of initial conditions. However, when the initial conditions are well approximated by a normal distribution with variance σ2, the denominator in the Boltzmann weight should be changed accordingly and this will change the final form of the first passage time distribution. Nonetheless, the derivation we present is general and can be adapted for a given p(s), see Appendix B.

[54] W. T. Coffey, Y. P. Kalmykov, and J. T. Waldron, The Langevin Equation, 2nd ed. (WORLD SCIENTIFIC, 2004).

[55] J. Desponds, H. Tran, T. Ferraro, T. Lucas, C. Perez Romero, A. Guillou, C. Fradin, M. Coppey, N. Dostatni, and A. M. Walczak, PLOS Computational Biology 12, 1 (2016).

[56] A. Cavagna, I. Giardina, and T. S. Grigera, Physics Reports 728, 1 (2018).

[57] A similar observation has been done in the analysis of spatial correlation functions in flocks of birds [118].

[58] D. Givon, R. Kupferman, and A. Stuart, Nonlinearity 17, 1 (2004).

[59] R. R. Coifman, I. G. Kevrekidis, S. Lafon, M. Maggioni, and B. Nadler, Multiscale Modeling & Simulation 7, 842 (2008), 10.1137/070696325.

[60] D. Giannakis, Applied and Computational Harmonic Analysis 47, 338 (2019).

[61] L. Callaham, J.-C. Loiseau, G. Rigas, and S. L. Brunton, Nonlinear stochastic modeling with langevin regression (2020), arXiv:2009.01006 [cond-mat.stat-mech].

[62] A. Frishman and P. Ronceray, Phys. Rev. X 10, 021009 (2020).

[63] F. Dietrich, A. Makeev, G. Kevrekidis, N. Evangelou, T. Bertalan, S. Reich, and I. G. Kevrekidis, Chaos: An Interdisciplinary Journal of Nonlinear Science 33, 023121 (2023).

[64] J. Berman, W. Bialek, and J. W. Shaevitz, Proceedings of the National Academy of Sciences 104, 20167 (2016).

[65] V. Alba, G. J. Berman, W. Bialek, and J. W. Shaevitz, Exploring a strongly non-markovian animal behavior (2020), arXiv:2012.15681 [q-bio.NC].

[66] E. Korobkova, T. Emonet, J. M. G. Vilar, T. S. Shimizu, and P. Cluzel, Nature 428, 574 (2004).

[67] O. Miramontes, O. DeSouza, L. R. Paiva, A. Marins, and S. Orozco, PLOS ONE 9, 1 (2014).

[68] K. Jung, H. Jang, J. D. Kralik, and J. Jeong, PLOS Computational Biology 10, 1 (2014).

[69] N. E. Humphries, N. Queiroz, J. R. M. Dyer, N. G. Pade, M. K. Musyl, K. M. Schaefer, D. W. Fuller, J. M. Brunnschweiler, T. K. Doyle, J. D. R. Houghton, G. C. Hays, C. S. Jones, L. R. Noble, V. J. Wearmouth, E. J. Southall, and D. W. Sims, Nature 465, 1066 (2010).

[70] D. W. Sims, E. J. Southall, N. E. Humphries, G. C. Hays, C. J. A. Bradshaw, J. W. Pitchford, A. James, M. Z. Ahmed, A. S. Brierley, M. A. Hindell, D. Morritt, M. K. Musyl, D. Righton, E. L. C. Shepard, V. J. Wearmouth, R. P. Wilson, M. J. Witt, and J. D. Metcalfe, Nature 451, 1098 (2008).

[71] D. A. Raichlen, B. M. Wood, A. D. Gordon, A. Z. P. Mabulla, F. W. Marlowe, and H. Pontzer, Proceedings of the National Academy of Sciences 111, 10.1073/pnas.1318616111 (2014).

[72] D. W. Sims, A. M. Reynolds, N. E. Humphries, E. J. Southall, V. J. Wearmouth, B. Metcalfe, and R. J. Twitchett, Proceedings of the National Academy of Sciences 111, 10.1073/pnas.1405966111 (2014).

[73] M. Viswanathan, S. V. Buldyrev, S. Havlin, M. G. da Luz, E. P. Raposo, and H. E. Stanley, Nature 401, 911 (1999).

[74] E. Wosniack, M. C. Santos, E. P. Raposo, G. M. Viswanathan, and M. G. E. da Luz, Phys. Rev. E 91, 052119 (2015).

[75] E. Wosniack, M. C. Santos, E. P. Raposo, G. M. Viswanathan, and M. G. E. da Luz, PLOS Computational Biology 13, 1 (2017).

[76] B. Guinard and A. Korman, Science Advances 7, eabe8211 (2021).

[77] A. Clementi, F. d’Amore, G. Giakkoupis, and E. Natale, in Proceedings of the 2021 ACM Symposium on Principles of Distributed Computing, PODC’21 (Association for Computing Machinery, New York, NY, USA, 2021) p. 81–91.

[78] G. H. Pyke, Methods in Ecology and Evolution 6, 1 (2015).

[79] A. Reynolds, Physics of Life Reviews 14, 59 (2015).

[80] Y. Tu and G. Grinstein, Phys. Rev. Lett. 94, 208101 (2005).

[81] T. S. Biró and A. Jakovác, Phys. Rev. Lett. 94, 132302 (2005).

[82] I. Lubashevsky, R. Friedrich, and A. Heuer, Phys. Rev. E 79, 011110 (2009).

[83] A. M. Reynolds and N. T. Ouellette, Scientific Reports 6, 30515 (2016).

[84] Interestingly, fast-fluctuating hopping rates and scale-invariance arguments have been used to explain heavy-tailed distributions of uncorrelated resting times in mice [119].

[85] D. G. Hernández, C. Rivera, J. Cande, B. Zhou, D. L. Stern, and G. J. Berman, eLife 10, e61806 (2021).

[86] S. Petrovskii, A. Mashanova, and V. A. A. Jansen, Proceedings of the National Academy of Sciences 108, 8704 (2011).

[87] W. Bialek and J. W. Shaevitz, Long time scales, individual differences, and scale invariance in animal behavior (2023), arXiv:2304.09608 [q-bio.NC].

[88] B. Qiao, C. Li, V. W. Allen, M. Shirasu-Hiza, and S. Syed, eLife 7, e34497 (2018).

[89] E. Overman, D. M. Choi, K. Leung, J. W. Shaevitz, and G. J. Berman, PLOS Computational Biology 18, 1 (2022).

[90] S. Wheatland, P. A. Sturrock, and J. M. McTiernan, The Astrophysical Journal 509, 448 (1998).

[91] G. Boffetta, V. Carbone, P. Giuliani, P. Veltri, and A. Vulpiani, Phys. Rev. Lett. 83, 4662 (1999).

[92] M. Beggs and D. Plenz, Journal of Neuroscience 23, 10.1523/JNEUROSCI.23-35-11167.2003 (2003).

[93] M. Newman, Contemporary Physics 46, 323 (2005), 10.1080/00107510500052444.

[94] J.-P. Bouchaud and A. Georges, Physics Reports 195, 127 (1990).

[95] D. ben Avraham and S. Havlin, Diffusion and Reactions in Fractals and Disordered Systems (Cambridge University Press, 2000).

[96] L. Cocchi, L. L. Gollo, A. Zalesky, and M. Breakspear, Progress in Neurobiology 158, 132 (2017).

[97] J. O’Byrne and K. Jerbi, Trends in Neurosciences 45, 820 (2022).

[98] T. Mora and W. Bialek, Journal of Statistical Physics 144, 268 (2011).

[99] F. den Hollander, Long time tails in physics and mathematics, in Probability and Phase Transition, edited by G. Grimmett (Springer Netherlands, Dordrecht, 1994) pp. 123–137.

[100] B. J. Alder and T. E. Wainwright, Phys. Rev. A 1, 18 (1970).

[101] J. Touboul and A. Destexhe, Phys. Rev. E 95, 012413 (2017).

[102] V. Priesemann and O. Shriki, PLOS Computational Biology 14, 1 (2018).

[103] M. Morrell, I. Nemenman, and A. J. Sederberg, Neural criticality from effective latent variable (2023).

[104] D. J. Schwab, I. Nemenman, and P. Mehta, Physical Review Letters 113, 068102 (2014).

[105] K. R. Ghusinga, J. J. Dennehy, and A. Singh, Proceedings of the National Academy of Sciences 114, 10.1073/pnas.1609012114 (2017).

[106] X. M. de Wit, A. van Kan, and A. Alexakis, Journal of Fluid Mechanics 939, R2 (2022).

[107] C. Costa and M. Vergassola, Fluctuating landscapes and heavy tails in animal behavior, 10.5281/zenodo.10030151 (2023).

[108] J. E. Sulston and S. Brenner, Genetics 77, 95 (1974).

[109] E. M. Bollt and N. Santitissadeekorn, Applied and computational measurable dynamics (Society for Industrial and Applied Mathematics, Philadelphia, United States, 2013).

[110] A. Bittracher, P. Koltai, S. Klus, R. Banisch, M. Dellnitz, and C. Schütte, Journal of Nonlinear Science 28, 471 (2018).

[111] L. McInnes, J. Healy, and J. Melville, Umap: Uniform manifold approximation and projection for dimension reduction (2018).

[112] V. A. Epanechnikov, Theory of Probability & Its Applications 14, 153 (1969), 10.1137/1114019.

[113] W. K. Härdle, M. Müller, S. Sperlich, and A. Werwatz, Nonparametric and Semiparametric Models (Springer Berlin, Heidelberg, 2006).

[114] W. Horsthemke and R. Lefever, Noise-Induced Transitions: Theory and Applications in Physics, Chemistry, and Biology (Springer Berlin, Heidelberg, 2006).

[115] We note that for the general dynamics of Eqs. (2,3), ω0 may have a s dependency. However, without loss of genserality, we consider that ω0 and ΔU (s) can be redefined to move the s dependency to the exponential as a subdominant contribution.

[116] R. Friedrich, J. Peinke, M. Sahimi, and M. Reza Rahimi Tabar, Physics Reports 506, 87 (2011).

[117] G. van Kampen, Journal of Statistical Physics 24, 175 (1981).

[118] A. Cavagna, A. Cimarelli, I. Giardina, G. Parisi, R. Santagati, F. Stefanini, and M. Viale, Proceedings of the National Academy of Sciences 107, 11865 (2010).

[119] A. Proekt, J. R. Banavar, A. Maritan, and D. W. Pfaff, Proceedings of the National Academy of Sciences 109, 10.1073/pnas.1206894109 (2012).

